# Engaging Visual Media Shifts Taste-Related Neural Processing: An fMRI Study on Distracted Eating

**DOI:** 10.64898/2026.02.20.706951

**Authors:** Robert Friedmann, Bichr Grii, Esther Jacoby, Ilya Digel, Rea Rodriguez-Raecke, Rik Sijben

## Abstract

**Background:** Distracted eating is prevalent in modern environments. While behavioral research consistently shows that distraction attenuates taste perception and increases food intake, the underlying neural mechanisms appear to be more complex.

**Objective:** This functional magnetic resonance imaging (fMRI) study investigated whether naturalistic distraction modulates gustatory processing via sensory suppression or reallocation of neural resources, as observed in more controlled cognitive load paradigms.

**Methods:** Thirty-eight healthy participants received sweet and umami taste stimuli of low and high concentration during fMRI scanning. Attentional state was manipulated using short food-related (low-distraction) versus film-related (high-distraction) video clips. After each video, participants rated perceived intensity and pleasantness. Group-level analyses included covariates for sex, body mass index (BMI), and hunger level.

**Results:** High distraction attenuated perceived intensity (p < 0.001, d = -0.28) and pleasantness (p < 0.01, d = -0.21), independent of taste category or concentration. No significant attenuation by distraction was observed in core gustatory regions (insula, orbitofrontal cortex). Instead, distraction increased activation in occipital, thalamic, and cerebellar regions, indicating a redistribution of processing resources toward visual and attentional systems.

**Conclusion:** Distraction reduced taste salience without lowering gustatory cortex activity, supporting resource-competition models rather than active sensory suppression. These results reinforce that the impact of distracted eating is behaviorally robust yet neurally subtle, highlighting the need for personalized stimuli and ecologically valid methods to capture real-world eating behavior. The study demonstrates that video-based paradigms work reliably in fMRI and capture how naturalistic distraction alters taste experience.

## 1 Introduction

Eating under distraction has become pervasive in modern environments, particularly among younger populations who frequently combine meals with digital media (Enes & Lucchini, 2016; Lissner et al., 2012; Utter et al., 2006). Distracted eating is consistently linked to increased food intake, reduced satiety, and weight gain (Avery et al., 2016; Ogden et al., 2013; Robinson et al., 2013; Shin, 2024). Although these behavioral effects are well established, much less is known about how distraction alters the perception of taste itself. An attenuation of perceived intensity or pleasantness may weaken satiation signals and facilitate overconsumption (Duif et al., 2020; Ruda, Chellapandian, & Freiherr, 2024), shaping eating habits in everyday life.

Recent evidence suggests that attentional state modulates taste perception. However, most existing studies rely on artificial cognitive load tasks (Liang et al., 2018; Razzaghi-Asl et al., 2024; Skorka-Brown et al., 2014). As a result, their ecological validity is limited, particularly given that real-world distracted eating typically involves naturalistic media, such as video viewing, which engages visual, affective, and attentional systems simultaneously and may interact with gustatory processing differently than controlled load tasks. A further limitation is that previous work has predominantly focused on sweet taste (Skałbania et al., 2025; van Meer et al., 2023; M. G. Veldhuizen et al., 2007), often omitting other taste modalities that are highly relevant to real-world meals, including umami (Kurihara, 2015), whose effects on appetite, self-control, and food choice remain only partially understood (Diepeveen et al., 2022; Magerowski et al., 2018).

### 1.1 Taste Perception

#### 1.1.1 General Gustatory Processing

Previous fMRI studies have consistently shown that gustatory stimulation reliably engages the anterior and middle insula, commonly referred to as primary gustatory cortex (Oliveira-Maia et al., 2011; M. G. Veldhuizen et al., 2007). Consistent with its role as a central relay for sensory information, thalamic involvement, particularly within the ventral posteromedial nucleus (VPM), has been reported during taste stimulation (Oliveira-Maia et al., 2011). These regions show robust BOLD responses across taste categories and concentrations, making them a central target for investigating neural correlates of taste perception. In contrast, posterior insula involvement appears less consistent and is often attributed to somatosensory or interoceptive processing rather than core gustatory functions (Brooks et al., 2005; Canna et al., 2019). Through extensive connections to salience, interoceptive, and reward networks, the insula further operates as a multimodal integrative hub (Gogolla, 2017; Menon, 2025; Small, 2010)

Beyond primary gustatory regions, the orbitofrontal cortex (OFC) has been shown to represent taste information by integrating gustatory input with hedonic evaluation and decision-related processes, supporting its role as secondary gustatory cortex (Bender et al., 2009; Duif et al., 2020; Small et al., 2007). OFC responses are particularly sensitive to changes in perceived pleasantness and motivational relevance, rather than sensory intensity alone.

Taste-related activity further extends into reward- and salience-related regions. For example, the amygdala has been shown to respond to emotionally salient food-related stimuli across modalities, with activation driven primarily by arousal and motivational relevance rather than valence alone (Bermudez-Rattoni, 2014; Small et al., 2003; van Dillen & van Steenbergen, 2018; Zald, 2003). Functional interactions between the insula and amygdala contribute to the affective tagging of taste memories and to contextual modulation under conditions such as distraction (J. O’Doherty et al., 2001). This is particularly relevant for umami, whose hedonic interpretation in Western populations has been shown to depend strongly on semantic and cultural context (Delwiche, 1996; Diepeveen et al., 2022).

Evidence regarding hemispheric asymmetries in insula function remains mixed, with reports of unilateral, bilateral, or attribute-specific activation patterns (e.g., intensity vs. pleasantness) (Bender et al., 2009; Dalenberg et al., 2015; Iannilli & Gudziol, 2019; Kurth et al., 2010), underscoring the need for cautious interpretation of lateralized BOLD responses.

#### 1.1.2 Attentional Modulation of Gustatory Processing

Experimental paradigms typically contrast conditions in which attention is directed toward the taste (“mindful eating”) (Grabenhorst & Rolls, 2008; van Rijn et al., 2018; M. G. Veldhuizen et al., 2007) with those in which it is diverted by external media or cognitive tasks (“distracted eating”) (Liguori et al., 2020; Ruda, Chellapandian, & Freiherr, 2024; van der Wal & van Dillen, 2013). Directing attention toward taste enhances activity in gustatory and reward regions, while focusing on different attributes (intensity vs. pleasantness) engages partially distinct circuits within the insula–OFC network (Grabenhorst & Rolls, 2008; van Rijn et al., 2018). Novel or ambiguous tastes evoke stronger insular activation, reflecting salience and sensory learning (Bermudez-Rattoni, 2014; Singh et al., 2015). Viewing food images alone can engage gustatory regions, demonstrating extensive cross-modal integration (Killgore et al., 2003; Simmons et al., 2005). Together, these findings reflect task-dependent top-down modulation of the gustatory cortex (Grabenhorst & Rolls, 2010; M. G. Veldhuizen et al., 2012).

Beyond selective attention, cognitive load introduces competition for limited attentional resources (Kahneman, 1973; Lavie et al., 2004). High load reliably reduces perceived taste intensity and weakens memory for recent eating episodes (Liang et al., 2018; van der Wal & van Dillen, 2013). In some studies, effects are more pronounced for high-intensity tastes, suggesting direct competition within a fixed-capacity gustatory system (Razzaghi-Asl et al., 2024; van Meer et al., 2023). Neuroimaging work demonstrates that load can attenuate insular responses or disrupt connectivity with valuation regions such as the OFC and nucleus accumbens, even when mean activation remains stable, indicating weakened integration of sensory and hedonic information (Duif et al., 2020; van Meer et al., 2023). Similar suppression effects in olfaction point to a modality-general mechanism of chemosensory downregulation under cognitive demand (Hoffmann-Hensel et al., 2017; Ruda, Chellapandian, Rott, et al., 2024).

### 1.2 Aims and Hypotheses

The present study aims to characterize how naturalistic distraction influences both sensory (intensity) and hedonic (pleasantness) aspects of taste perception across sweet and umami stimuli using an ecologically valid video-based paradigm combined with event-related fMRI. Assessing pleasantness represents a key conceptual extension, as hedonic experience is central to food choice (Kringelbach, 2004) but has been largely neglected in prior neuroimaging work. The guiding question is therefore not only whether distraction affects how much we eat, but whether it changes how food tastes.

At the behavioral level, distraction was expected to reduce perceived taste intensity and pleasantness, consistent with prior load-related attenuation effects (Liang et al., 2018; van der Wal & van Dillen, 2013). At the neural level, we anticipated that suppression in the anterior insula and OFC would be moderate and less pronounced than the effects reported in artificial load tasks (Duif et al., 2020; Skałbania et al., 2025) — alongside reduced involvement of reward-related regions and increased activation in visual and attentional networks during high distraction. Finally, stronger distraction effects were expected for high-concentration stimuli due to increased resource competition (Razzaghi-Asl et al., 2024; van Meer et al., 2023), whereas no substantial interaction with taste category (sweet vs. umami) was predicted.

## 2 Methods

### 2.1 Participants

Forty-two healthy volunteers aged 18 to 34 participated in the study (Median age = 23 years, IQR = 21–25; BMI: 23.0 ± 2.9 kg/m^2^; male = 20, female = 22). Biological sex rather than gender is reported in accordance with study documentation. Participants were recruited through university buildings and student residences in Aachen and Jülich and through the University Hospital RWTH Aachen. Eligibility required an age of 18 to 40, a BMI within the extended normal range according to WHO criteria (17.0 – 29.9 kg/m^2^), no neurological or psychiatric disorders, no metabolic disease (e.g. diabetes), non-smoking status, no susceptibility to motion sickness, and no contraindications for MRI or the administered taste stimuli. The BMI restriction avoided potential taste alterations associated with obesity (Ruda, Chellapandian, & Freiherr, 2024; van Meer et al., 2022), and smokers were excluded due to known effects of smoking on taste (Grunberg, 1982; Kale et al., 2019). Individuals prone to motion sickness were excluded because two of the videos presented during the experiment showed roller coaster rides. All participants completed the experiment in either German or English, according to their preference.

Prior to the measurement day, participants received safety information and a description of the task. To control for hunger level, participants were instructed to abstain from eating for the last three hours and to drink only water in the last hour preceding the measurement. On the measurement day, participants gave written consent and all MRI safety procedures were explained. Participants unfamiliar with umami received a brief explanation of the term. A short pre-experiment questionnaire ensured naive status regarding the study purpose, and a short post-experiment questionnaire was completed after scanning.

The study conformed to the ethical principles of the World Medical Association’s Declaration of Helsinki. The study was approved by the Ethics Committee of the Medical Faculty of RWTH Aachen University (EK 24-242) and was pre-registered in the German Clinical Trials Register (DRKS00034003) of the German Federal Institute for Drugs and Medical Devices (BfArM).

### 2.2 Procedure

Participants performed two consecutive sessions of a distraction task while undergoing fMRI scanning. The task consisted of watching muted videos and answering questions via keyboard input after each video. During video presentation, liquid taste stimuli were administered intraorally via a multi-channel gustometer. An overview is shown in Figure 1.

**Figure 1:**
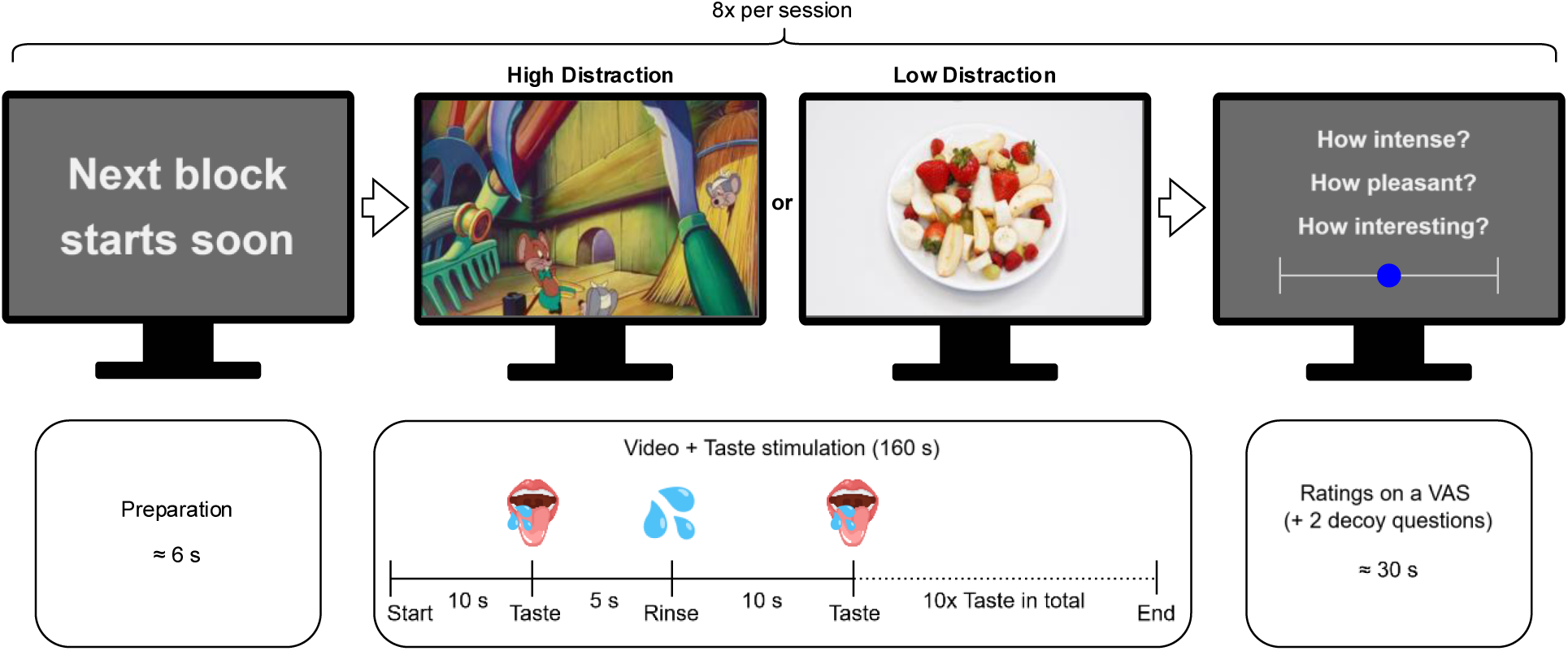
Depiction of one block of the paradigm. The experiment contained two consecutive sessions, each comprising eight blocks, covering all conditions of the 2×2×2 within-subject factorial design. A short preparation screen separated the blocks. Each block consisted of a 160-second video (high or low Distraction). Starting 10 seconds after video onset, a taste stimulus was administered every 15 seconds, followed by a rinse 5 seconds later. After each video, participants rated the intensity and pleasantness of the taste and their interest in the video.

The experiment followed a 2×2×2 factorial design with *Distraction* (high vs. low), *Taste* (sweet vs. umami) and *Concentration* (high vs. low) as within-subject factors. Each session comprised eight blocks, one for each possible combination of these factors. Each block consisted of a 160-second video (high or low distraction) and the corresponding taste stimulus (sweet or umami at high or low concentration). Block order was pseudorandomized and identical across participants within a session but differed between sessions. The exact sequence is shown in Supplementary Figure 1.

Before the first block, we instructed participants to watch the videos and swallow the liquids when delivered. A six-second countdown preceded each block. After each block, a brief reminder screen appeared for five seconds, followed by five questions presented as unanchored visual analogue scales (VAS). Participants rated perceived *intensity* and *pleasantness* of the associated taste stimuli, rated the associated video’s *interest* level, and answered two masking questions about viewing frequency and taste–video matching.

Before the first session, participants completed a three-minute practice run to familiarize themselves with the controls and with the umami stimulus. A structural scan was acquired during a five-minute break between the two sessions.

### 2.3 Gustatory Stimulation

The sweet stimuli consisted of low- and high-concentration sucrose solutions with intended concentrations of 5 wt% (0.15 M) and 15 wt% (0.44 M). The umami stimuli consisted of low- and high-concentration monosodium glutamate (MSG) solutions with intended concentrations of 0.5 wt% (0.03 M) and 2.5 wt% (0.15 M). The solutions were prepared freshly before each measurement, kept at room temperature (21–23 °C), and the actual concentrations were 5.00 ± 0.012 wt% and 14.97 ± 0.071 wt% for sucrose and 0.50 ± 0.002 wt% and 2.50 ± 0.002 wt% for MSG. Concentrations were chosen based on pilot testing to be clearly distinguishable within each taste and approximately isointense across tastes.

All taste solutions and the rinse were prepared using still bottled water of the same brand (Aldi Quellbrunn Naturell). Each taste stimulus and each rinse had a volume of 2.61 ± 0.05 mL. During each block, ten taste stimuli were delivered at intervals of fifteen seconds beginning ten seconds after video onset. A rinse followed each stimulus five seconds later. The final rinse occurred at 150 seconds (Figure 1).

In each block, eight of the ten stimuli matched the taste category and concentration of that block (target stimuli). The remaining two were distractor stimuli of the opposite taste to reduce habituation. Their positions were pseudorandomized, identical across participants, and never occurred first, last, or consecutively. Supplementary Table 1 provides the exact presentation order.

### 2.4 Visual Distraction

The high-distraction condition included four entertainment-related clips covering an action scene, a silent film, a cartoon and a roller coaster ride. In the second session, the continuations of the clips from the first session were shown, except for the roller coaster, which featured a different ride. These videos were selected based on pilot interest ratings and were either royalty-free or used with permission. High-distraction videos were paired pseudorandomly with the taste and concentration conditions, with the pairing kept consistent across participants.

The low-distraction videos were designed to resemble an eating context and showed dishes being consumed in motion rather than static images. Each clip displayed a sweet or savory dish on a white background while a hand entered the frame periodically to remove a portion of the food. Slight screen shake and minimal camera motion were added during editing to approximate natural head movement. Two sweet and two savory dishes were shown per session and were congruently paired with the corresponding sweet or umami taste conditions. Different dishes were used in the second session while maintaining congruent pairing. All low-distraction videos were filmed and edited at the Audiovisual Media Center of the Medical Faculty of RWTH Aachen University.

All videos were adapted to a 16:10 aspect ratio to standardize visual presentation and were shown without sound to avoid auditory activation. Representative screenshots are provided in Supplementary Figure 1.

### 2.5 Devices

Participants viewed the task on an MRI-compatible monitor (BOLDscreen, 1920 × 1200 pixels, Cambridge Research Systems Ltd, Rochester, England) positioned behind the scanner bore and reflected via an angled mirror above the eyes. Responses were recorded with a custom optical keypad placed under the right hand with left, confirm and right buttons. To minimize scanner noise and avoid auditory activation, participants wore earplugs and OptoActive-II active noise canceling headphones (Optoacoustics Ltd., Israel).

The experiment was programmed in PsychoPy 2023.2.3 (Peirce et al., 2019). Stimulus delivery was controlled directly through PsychoPy, which triggered the gustometer (ETT Gustometer 2+, Emerging Tech Trans LLC, College Station, TX, USA) to dispense liquids through peristaltic pumps (Figure 2). For each stimulus, PsychoPy sent a trigger activating the corresponding pump for 375 milliseconds. The gustometer was located in the control room, and taste solutions were transported through eight-meter tubing routed into the scanner room. Five channels were used, corresponding to four taste solutions and one rinse, each assigned to a single liquid for the duration of the study to prevent cross-contamination. All channels converged in a pacifier-like mouthpiece with separate outlets.

**Figure 2:**
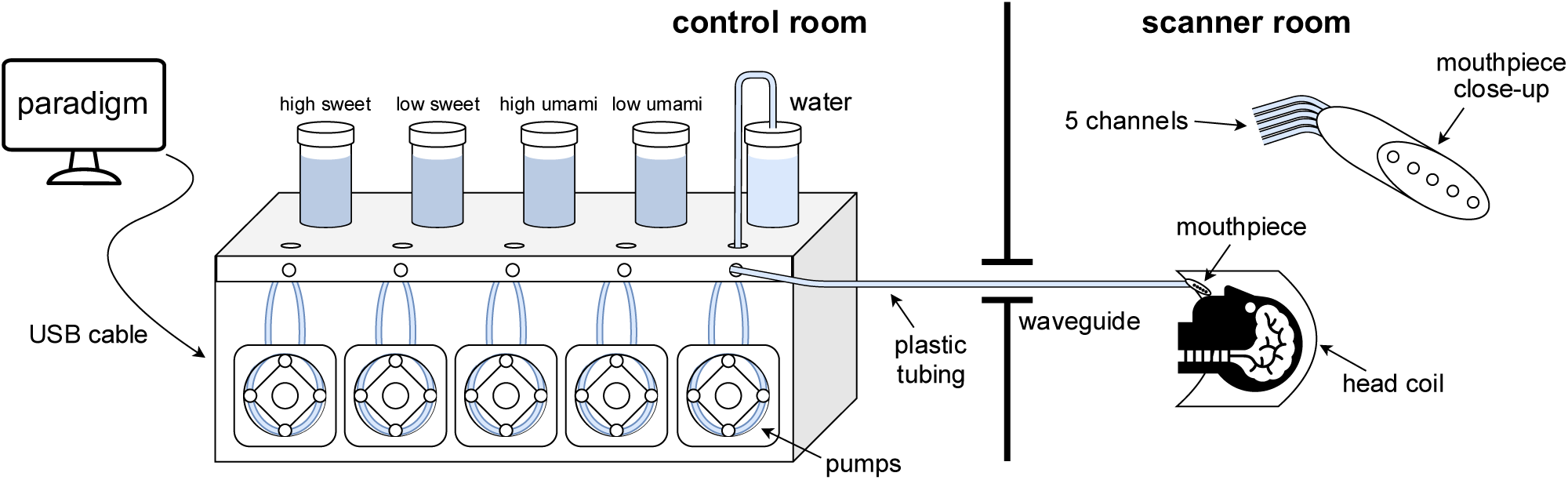
Depiction of the multi-channel gustometer. The device was connected to a computer running the paradigm and controlling the gustometer’s peristaltic pumps, triggering stimulus delivery. Each tastant was assigned a channel for the duration of the study. The solutions were routed from glass containers through plastic tubing, via the waveguide and an 8-m-long tube into the scanner room. The end of the tubing was fitted with a mouthpiece that the participants held in their mouth during the experiment.

The mouthpiece was fixed to an articulating plastic arm mounted on the head coil and positioned in the participant’s mouth, allowing stimuli to be delivered directly to the tongue while minimizing swallowing-induced head and jaw motion. Before each measurement, the system was primed by flushing all channels simultaneously, followed by individual testing to ensure stable flow.

### 2.6 MRI Data Acquisition

The study was conducted on a 3 T Siemens Magnetom Prisma scanner (syngo MR XA60 platform). The images were acquired using a 20-channel head coil. During the two paradigm sessions, functional images were acquired using a T2*-weighted Gradient Echo (GRE) Echo-Planar Imaging (EPI) sequence with the following parameters: TR = 2020 ms, TE = 37 ms, FA = 72°, FOV = 220 mm, 72 transverse slices, voxel size = 2.0 mm isotropic. The sequence was multi-band accelerated with a factor of 3. The mean number of volumes in a functional sequence was 785 with a mean total acquisition time of 26 minutes and 25 seconds per run.

Between the two paradigm sessions, a high-resolution structural image was acquired using a T1-weighted Magnetization Prepared Rapid Gradient Echo (MPRAGE) sequence with the following parameters: TR = 2300 ms, TE = 2.32 ms, FA = 8°, FOV = 240 mm, 192 sagittal slices, voxel size = 0.9 mm isotropic. The total acquisition time was 5 minutes and 21 seconds.

The B0 field was mapped using a GRE field map with the following parameters: TR = 438 ms, TE1 = 4.92 ms, TE2 = 7.38 ms, FA = 60°, FOV = 228 mm, 45 slices, voxel size = 3.0 mm isotropic.

### 2.7 Quality Control and Manipulation Check

Data from two participants were excluded due to structural brain anomalies. Data extraction and quality control were performed in MATLAB R2023b (The MathWorks, Inc., 2023). Two additional participants had incomplete PsychoPy logfiles, resulting in five missing data points (out of 640 total data points across the study). Potential gustometer failures were screened by identifying very low intensity ratings combined with neutral pleasantness ratings, yet no data points met the exclusion criteria.

Validation of the experimental conditions was conducted in R version 4.5.0 (R Core Team, 2021). Tests of homoscedasticity indicated equal variances across sexes for intensity ratings but not for pleasantness ratings, therefore sex was included as a fixed effect in the pleasantness models. Plots of intensity and pleasantness ratings across Concentration and Taste revealed strong interindividual variability in taste preference patterns (Supplementary Figure 2), motivating the inclusion of a random slope for Taste in the mixed models.

To confirm the distinctiveness of the stimuli, paired t-tests showed robust differences between high and low Concentration within each taste (both p < 0.001) and no significant differences across tastes (p_high_ = 0.465, p_low_ = 0.237), indicating that sweet and umami stimuli were comparable in baseline intensity (Supplementary Figure 3). The distraction manipulation was validated using the interest ratings, which were significantly higher for the high-distraction videos with a large effect size (p < 0.001, d = 1.31) (Supplementary Figure 3).

Paired t-tests across sessions revealed no significant differences in any rating measure (all p > 0.1), therefore session was not included as a factor in the subsequent behavioral models.

### 2.8 Behavioral Analysis

Behavioral analyses were conducted in R version 4.5.0 using linear mixed-effects models fitted with lmer (lme4 package). *Intensity Rating* and *Pleasantness Rating* were modeled separately as dependent variables. Model selection followed likelihood-ratio tests and information criteria, while retaining predictors relevant to the study hypotheses.

*Concentration*, *Taste* and *Distraction* were considered as fixed effects in both models. *Sex* was considered for the pleasantness model due to variance differences identified during quality control. *BMI* and *Hunger Level* were considered in both models. Both models included a random intercept for *Participant* and a random slope for Taste, based on the quality control analyses indicating strong interindividual variation along this dimension. Across all candidate models, the addition of a random slope for Taste consistently produced the strongest improvement in model fit. All fixed effects were included additively unless interactions improved model fit. Main effects were retained whenever an associated interaction was present to ensure proper interpretation of the model parameters.

For the intensity model, none of the interaction terms improved model fit. The final model was:

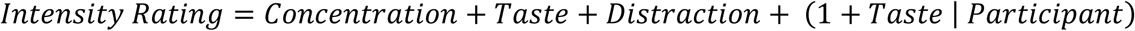

The pleasantness model was improved by several effects and interactions. Its final structure was:

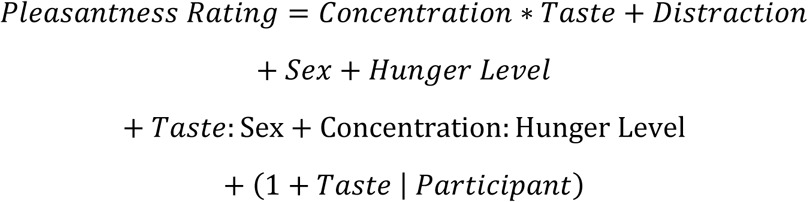

Model assumptions were evaluated using the performance package (Lüdecke et al., 2021), Shapiro– Wilk normality tests and skewness and kurtosis metrics. All checks were satisfactory and confirmed adherence to parametric and non-parametric test assumptions. Fixed effects were assessed with Type III ANOVA and significant effects were followed by post-hoc tests of estimated marginal means. Effect sizes were quantified as Cohen’s d and standardized coefficients were reported to indicate the relative influence of each predictor.

### 2.6 fMRI Analysis

#### 2.9.1 Preprocessing

Data from the same two participants excluded during the behavioral analysis were also excluded from the fMRI analysis due to structural anomalies. For two additional participants, one experimental session had to be excluded because missing event timing information in session 1 prevented valid modeling.

Preprocessing and all subsequent fMRI analyses were carried out in SPM12 (Eickhoff et al., 2005) running on MATLAB R2023b. Before preprocessing, all functional volumes were visually inspected in axial, sagittal and coronal views to verify proper coverage and identify artifacts. For one participant, a wrap-around artifact displaced frontal tissue. The affected volumes were corrected using a circular shift in MATLAB, and the missing two-pixel strip at the border of each slice was reconstructed by weighted interpolation of adjacent columns. No substantial positional inconsistencies were found when comparing sessions for each participant.

The first five volumes of each run were discarded to allow for signal stabilization. Field maps were used to generate voxel displacement maps which were applied during realignment and unwarping to correct magnetic field inhomogeneity. Functional images were then realigned to correct for motion. Each participant’s mean functional image was coregistered to their structural T1 image, and the anatomical image was normalized to MNI-152 space using SPM12 tissue probability maps. The resulting deformation fields were applied to the functional images, which were resampled to 2 mm isotropic voxels and smoothed with an 8 mm FWHM Gaussian kernel. Preprocessing quality was visually inspected at each step.

Motion traces in all six degrees of freedom were plotted and inspected (Supplementary Figure 4). Two participants exhibited pronounced motion spikes exceeding 2 mm translation and 2 degrees rotation and were excluded from further analyses. The thresholds of 2 mm translation and 2° rotation were chosen because, given the 2 mm spatial resolution, larger displacements would result in undesirable shifts within the regions of interest. This yielded a final sample of 36 complete and two partial datasets for fMRI analysis.

#### 2.9.2 Single-Subject Analysis

First-level analyses were conducted using a general linear model. The design matrix included eight regressors corresponding to the conditions of the 2×2×2 within-subject factorial design, with event onsets representing only the target stimuli of each condition. All rinse onsets were combined into one regressor that served as a control condition. Additional regressors modeled distractor taste events and pre-block and post-block periods to ensure that signal variance associated with these events was not misattributed. An implicit baseline, represented by the model intercept, accounted for signal not captured by the other regressors.

All events were modeled as zero-duration impulses and convolved with the canonical hemodynamic response function. Six motion parameters from realignment were included as nuisance regressors. A high-pass filter of 512 seconds removed low-frequency drifts while preserving signal related to the extended video presentation.

All main effects and interaction contrasts were computed at first level. Modeling interactions at first versus second level produced equivalent results. For participants missing one session, contrasts were computed from the available session, as both sessions contained the same conditions and design structure.

#### 2.9.3 Group Analysis

For each first-level contrast, a separate second-level model was created using a one-sample t-test. Sex, BMI and Hunger Level were included as mean-centered covariates to control for variability related to demographic or physiological factors, mirroring the behavioral models. Each model contained a positive and a negative contrast for the experimental effect, as well as covariate contrasts. The covariate contrasts were inspected for completeness but are not discussed further because none showed interpretable effects.

#### 2.9.4 ROI Analysis

A region of interest (ROI) analysis was conducted to examine distraction effects within key taste-related regions. This approach was chosen because, given the naturalistic paradigm, effects in the insula might be subtle or absent at the whole-brain level. Focusing on relevant subregions of the insula increases statistical power, allowing small differences to be detected and reducing the likelihood that null results reflect Type II errors rather than true absence of effect. ROIs were functionally defined using the group-level Stimulus > Rinse contrast, which captured general taste processing independent of distraction. Because this contrast was orthogonal to the distraction manipulation, the approach did not introduce circularity.

The insula was defined a priori as the primary ROI due to its significance in taste representation. It was represented by two separate significant clusters identified in the localizer contrast, one in the left anterior insula and one in the left middle insula. Both ROIs were extracted using the same threshold as the whole-brain analysis and exported as binary masks in MNI-152 space.

For both ROIs, mean β values were extracted from the first-level contrast representing the main effect of distraction (High Distraction > Low Distraction). This contrast was chosen because the behavioral data did not suggest any meaningful interactions with taste or concentration, making the main distraction contrast the most informative test of the hypothesis.

Group-level analyses of the extracted mean β values were performed in R. A one-sample t-test assessed whether ROI activation differed between high and low distraction. To maintain consistency with the whole-brain analyses, multiple regression models including Sex, BMI and Hunger Level were additionally fitted to evaluate whether these covariates explained further variance. Because the insula was specified a priori as a ROI, no correction for multiple comparisons was applied.

## 3 Results

### 3.1 Intensity Model

The intensity model included Concentration, Taste and Distraction as fixed effects with a random intercept and a random slope for Taste by Participant. Type III ANOVA indicated a strong effect of Concentration (F(1, 551.26) = 203.33, p < 0.001) and Distraction (F(1, 551.56) = 12.26, p < 0.001). Taste showed no significant effect (F(1, 39.02) = 1.98, p = 0.167) (Figure 3).

**Figure 3:**
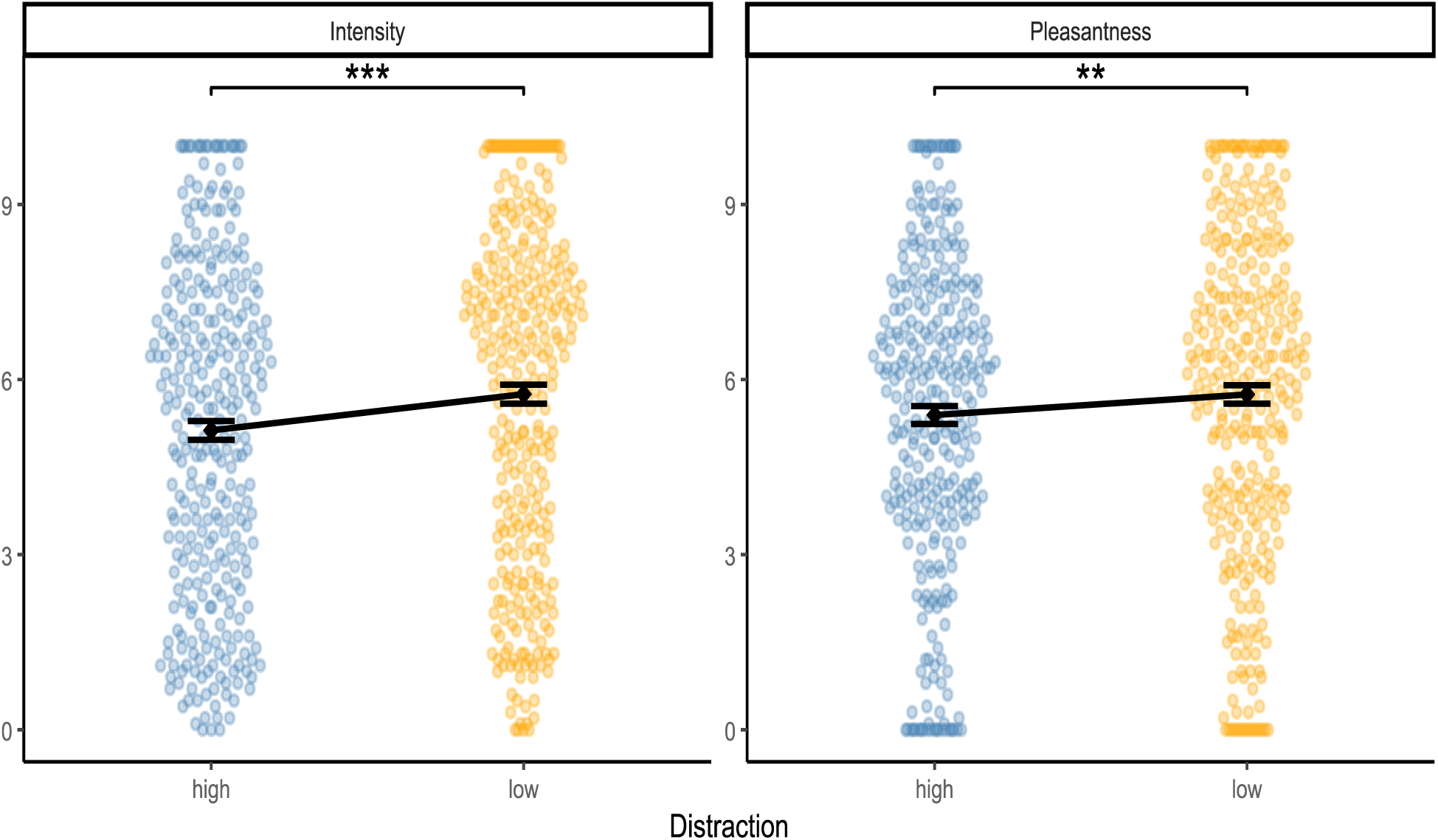
Effects of distraction on (left) intensity and (right) pleasantness. High distraction significantly decreased intensity and, to a lesser extent, pleasantness. Dots represent individual ratings; vertical lines represent standard error. ** indicates p < 0.01, *** indicates p < 0.001.

Post-hoc comparisons confirmed that high Concentration increased intensity ratings with a large effect size (estimate = 2.57, SE = 0.18, p < 0.001, d = 1.13). High Distraction caused a small decrease in intensity ratings (estimate = -0.63, SE = 0.18, p < 0.001, d = -0.28).

Standardized parameter estimates indicated that Concentration (high) had the strongest influence (std. β = 0.88), followed by Distraction (high) (std. β = -0.22), while Taste (umami) had a minimal influence (std. β = -0.13).

### 3.2 Pleasantness Model

The pleasantness model included Concentration, Taste, Distraction, Sex and Hunger Level as fixed effects with the interactions Taste × Sex, Concentration × Taste and Concentration × Hunger Level, and a random intercept and a random slope for Taste by Participant. Type III ANOVA showed significant main effects of Concentration (F(1, 548.99) = 10.07, p = 0.002), Taste (F(1, 38.06) = 55.15, p < 0.001), Distraction (F(1, 549.35) = 6.85, p = 0.009) and Sex (F(1, 36.94) = 4.83, p = 0.034). Hunger Level did not reach significance as a main effect (F(1, 37.06) = 2.03, p = 0.163). Significant interactions were observed for Taste × Sex (F(1, 38.06) = 14.59, p < 0.001), Concentration × Taste (F(1, 549.01) = 39.42, p < 0.001), and Concentration × Hunger Level (F(1, 548.98) = 8.85, p = 0.003) (Figure 4).

**Figure 4:**
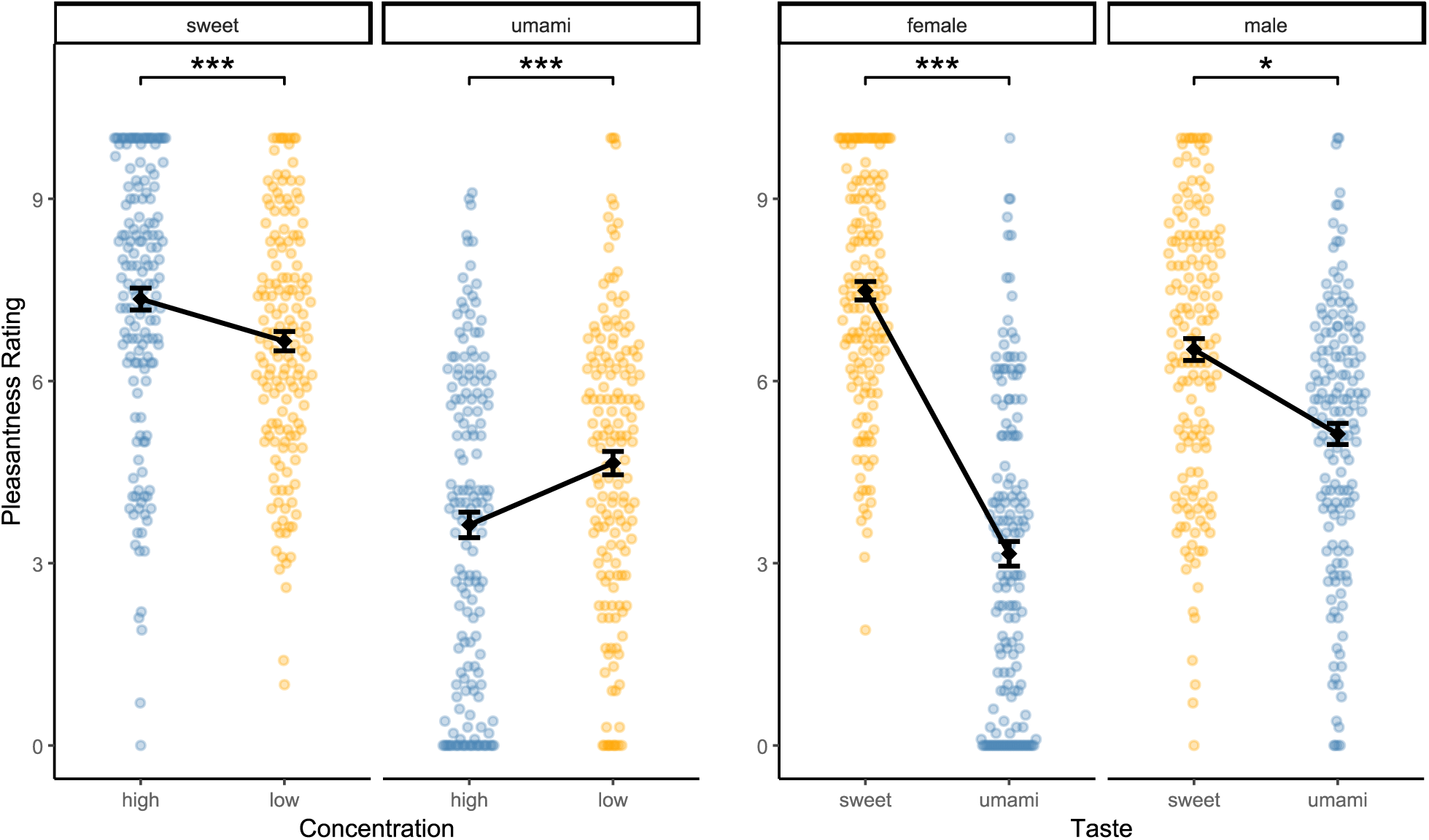
(Left) Effects of concentration on pleasantness for both tastes. High concentration significantly increased pleasantness for sweet but decreased it for umami. (Right) Effects of taste on pleasantness for both sexes. Female participants rated sweet substantially higher than umami, while male participants showed a smaller, yet significant, difference. Dots represent individual ratings; vertical lines represent standard error. * indicates p < 0.05, *** indicates p < 0.001.

Post-hoc comparisons revealed that high Concentration increased pleasantness ratings for sweet (estimate = 0.70, SE = 0.19, p < 0.001, d = 0.41) but decreased pleasantness ratings for umami (estimate = -0.998, SE = 0.19, p < 0.001, d = -0.59). The effect of Taste was moderated by Sex: Female participants rated sweet substantially higher than umami (estimate = 4.34, SE = 0.55, p < 0.001, d = 2.56), while male participants showed a smaller, yet significant, difference (estimate = 1.39, SE = 0.55, p = 0.015, d = 0.82). High Distraction caused a small reduction in pleasantness ratings (estimate = - 0.35, SE = 0.14, p = 0.009, d = -0.21).

Standardized parameter estimates indicated that Taste (umami) had the strongest negative influence (std. β = -1.87). The Taste × Sex (std. β = 1.07) and Concentration × Taste (std. β = 0.61) interactions also showed substantial effects. Concentration (high) and Sex (male) showed smaller negative influences (std. β = -0.25, std. β = -0.29). Distraction had a modest attenuating effect (std. β = -0.13). Hunger Level showed no meaningful effect, whereas the Concentration × Hunger Level interaction made a small contribution (std. β = -0.14).

### 3.3 fMRI Results

Whole-brain results were thresholded at p < 0.05, family wise error (FWE) corrected for multiple comparisons. For the concentration and interaction contrasts, where smaller effects were expected, we additionally inspected results at p < 0.001 combined with cluster-level FWE correction (p_FWEc_ < 0.05) to increase sensitivity. Anatomical labels were assigned using the Harvard–Oxford cortical and subcortical atlases and the probabilistic cerebellar atlas in MNI-152 space. None of the covariates (Sex, BMI, Hunger Level) showed significant effects at p_FWE_ < 0.05. Before assessing distraction-related effects, we verified that the taste stimuli elicited the expected engagement of gustatory and somatosensory regions. All active clusters are listed in Supplementary Tables 2–4.

#### 3.3.1 Main Effects of the Taste Stimuli

The contrast Stimulus > Rinse (with “Stimulus” comprising all taste events across levels, including distractor taste stimuli) produced a widespread activation pattern across occipital, parietal and subcortical regions, reflecting visual processing, swallowing and gustatory input. Importantly, taste delivery engaged the left anterior insula and adjacent middle insula, together with strong thalamic responses consistent with VPM involvement and gustatory regions in the brainstem. No right anterior insula activation was detected. Limbic engagement was present in the amygdalae. No regions responded more strongly to the rinse condition.

Comparing high to low concentration (at p_FWEc_ < 0.05) revealed strong activation in occipital regions and enhanced responses in the anterior and middle insula and thalamus. These effects demonstrate that our stimuli reliably recruited primary gustatory regions, with stronger insular responses at higher concentration, although insular activation remained predominantly left-hemispheric. No regions showed stronger activation for low concentration.

Taste category produced no differential activation in gustatory cortex. Contrasting sweet against umami showed differences only in occipital and temporal regions. Neither contrast revealed insular or OFC effects.

Together, these taste-related contrasts demonstrate robust activation in somatosensory, insular, occipital, and subcortical regions, with T-values indicating strong peak responses. These baseline patterns provide the functional context for evaluating the subsequent distraction-related contrasts.

#### 3.3.2 Distraction-Related Effects

##### 3.3.2.1 Main Effect of Distraction

High minus low distraction elicited a widespread increase in activation across visual and parietal regions, including strong engagement of occipital cortex, lingual gyrus and precuneus. These effects reflected the high visual and attentional demands of the distraction videos. Subcortically, distraction increased activation in thalamus bilaterally, alongside enhanced cerebellar responses. Additional activation was present in temporal areas typically involved in attentional processing. No attenuation was observed in gustatory regions. The inverse contrast revealed only small bilateral clusters in the lateral occipital cortex (Figure 5).

**Figure 5:**
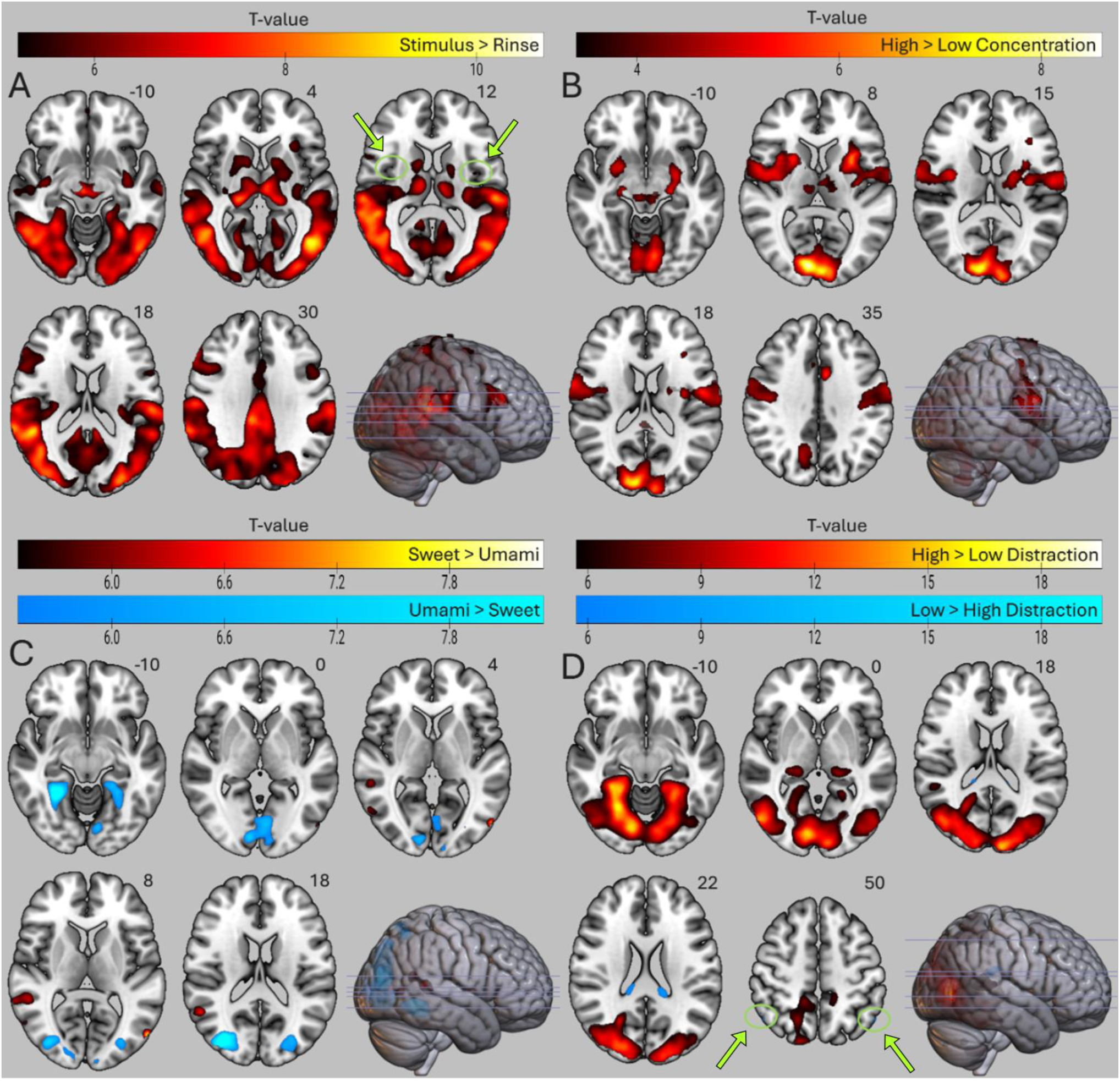
Visualization of the main effects of (A) Taste stimulus, (B) Concentration, (C) Taste category and (D) Distraction. (A) Stimulus > Rinse comprised all taste events and showed widespread activation in occipital and parietal cortex, subcortical structures, left anterior insula, and bilateral middle insula (green arrows); no right anterior insula. (B) High Concentration > Low Concentration revealed robust, bilateral insular and thalamic activation; no Low > High effects. (C) Sweet activated right angular gyrus, while Umami engages lateral occipital and fusiform regions. (D) High distraction elicited a large bilateral occipital cluster, pronounced lingual gyrus, left VPM, and cerebellar lobules; Low > High revealed only two small bilateral clusters in the superior lateral occipital cortex/angular gyrus (green arrows), and no gustatory activation. A, C, D significant at pFWE < 0.05; B significant at pFWEc < 0.05 to improve sensitivity. Left and right are indicated according to radiological convention.

##### 3.3.2.2 Distraction Effects by Concentration

The interaction between Distraction and Concentration (at p_FWEc_ < 0.05) showed that distraction effects were stronger in occipital cortex at high concentration, whereas at low concentration the effects were more pronounced in inferior temporal and lateral occipital areas. No interaction effects were observed in gustatory regions (Figure 6).

**Figure 6:**
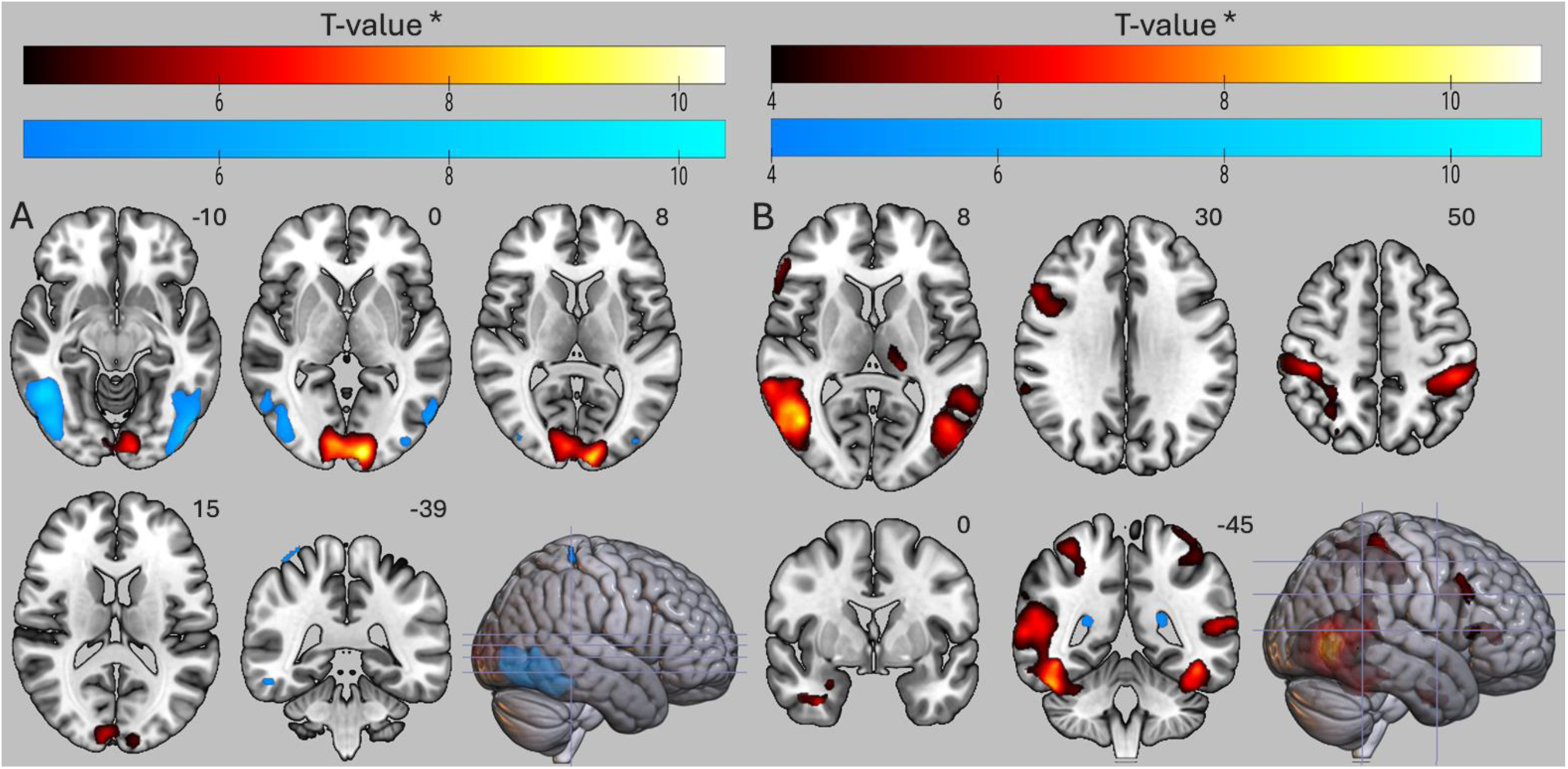
Visualization of the interaction effects. (A) Distraction (High > Low) × Concentration (High > Low) in red and Distraction (Low > High) × Concentration (High > Low) in blue. (B) Distraction (High > Low) × Taste (Sweet > Umami) in red and Distraction (Low > High) × Taste (Sweet > Umami) in blue. Both interaction contrasts showed predominantly bilateral activation in occipital regions, while the interaction with Taste revealed more widespread differences. Notably, no insular activation was observed (see axial slices 8 and 15). All significant at pFWEc < 0.05 to improve sensitivity. Left and right are indicated according to radiological convention.

When examining the simple effects, high distraction at high concentration produced a widespread visual–parietal activation pattern that extended across occipital cortex, lingual gyrus and precuneus, accompanied by bilateral thalamic engagement and cerebellar responses (Supplementary Table 4 A). These findings indicate that distraction strongly recruited visual and subcortical processing when taste intensity was high. No regions showed reduced activation under high distraction.

At low concentration, high distraction elicited a similar but weaker pattern centered on lateral occipital and inferior temporal regions (Supplementary Table 4 B). In contrast to the high-concentration condition, no thalamic engagement was observed, and cerebellar involvement was limited. Again, no regions showed attenuation under high distraction.

##### 3.3.2.3 Distraction Effects by Taste

The interaction between Distraction and Taste (at p_FWEc_ < 0.05) showed that distraction effects were stronger for sweet than for umami. For sweet, distraction produced larger differences in bilateral inferior lateral occipital cortex with extensions into middle and inferior temporal regions. Additional bilateral activation appeared in postcentral and superior parietal areas, and at a more liberal threshold the interaction included left thalamic involvement. No corresponding effects were observed for umami, and no interaction effects emerged in insula or OFC (Figure 6).

When examining the simple effects, high distraction during sweet taste produced a robust bilateral visual activation pattern involving occipital cortex and lingual gyrus, accompanied by bilateral thalamic engagement and additional responses in amygdala and precentral regions (Supplementary Table 4 C). No regions showed reduced activation under high distraction.

High distraction during umami taste produced a similar but attenuated pattern (Supplementary Table 4 D). Activation was centered on lingual and occipital regions with bilateral cerebellar involvement, but thalamic engagement was not present. Postcentral activation was observed bilaterally, and no attenuation under high distraction was detected.

#### 3.3.3 ROI Analysis of the Main Effect of Distraction

Mean parameter estimates from the High Distraction > Low Distraction contrast were extracted for the left anterior insula and the left middle insula. Each ROI was tested against zero using one-sample t-tests, with additional multiple regression models including Sex, BMI, and Hunger Level to assess covariate effects.

In the left anterior insula, activation did not differ significantly between high and low distraction (t(37) = -1.64, p = 0.11). The regression model did not improve explanatory power (F(3,34) = 1.80, p = 0.17), although a small negative association with Sex emerged (std. β = -3.46, p = 0.037), indicating slightly lower activation in male participants. The left middle insula showed no significant effect of distraction (t(37) = -1.33, p = 0.19) and no covariate effects (F(3,34) = 0.97, p = 0.42). Overall, the ROI analysis provided no evidence of distraction-related attenuation in the insula.

## 4 Discussion

The present study investigated how video-based distraction affects both sensory (intensity) and hedonic (pleasantness) aspects of taste perception across sweet and umami stimuli. We hypothesized that high distraction would reduce perceived intensity and pleasantness, particularly for high-concentration stimuli, while neural effects in primary and secondary gustatory regions would be moderate, reflecting the naturalistic mode of the distraction. We further anticipated increased engagement of visual and attentional networks under distraction, with no strong interactions with taste category.

### 4.1 Behavioral and Neural Responses to Taste

#### 4.1.1 Behavioral Taste Effects

Participants reliably rated higher concentrations as more intense across both tastes. Pleasantness, however, showed a clear Concentration × Taste interaction. Increasing sucrose concentration increased pleasantness, whereas concentrated MSG reduced it. This likely reflects differences in hedonic baselines and familiarity. Sucrose is inherently pleasant and the high-concentration solution (15 wt%) was comparable to common beverages such as Coca-Cola or apple juice (∼10 wt%), chocolate milk (∼11 wt%), and energy drinks (∼12 wt%). MSG can be palatable in moderate doses but becomes aversive when presented as a high-concentration liquid (Moldovan et al., 2021). Only 13 participants reported prior experience with pure MSG, which likely increased variability.

The sensory-reduced nature of the stimuli (no texture or retronasal olfaction) may have lowered the motivational salience of umami (Moldovan et al., 2021), while sweet taste retained strong learned reward associations (Ventura & Mannella, 2011). Alternating sweet and umami blocks may have further amplified contrast effects, reducing the relative hedonic value of umami.

A strong Taste × Sex interaction emerged for pleasantness, driven by higher sweet liking in female participants. Although larger than typically reported, the direction is consistent with prior reports of sex differences in sweet preference (Ruda, Chellapandian, & Freiherr, 2024; van Langeveld et al., 2018; Zellner, 1999). However, interindividual variability exceeded the contribution of sex. This indicates that previously reported sex differences may be inflated when interindividual variability is not modeled, a point also emphasized in recent work (Duif et al., 2020; van Meer et al., 2023). The substantial participant-level variance observed here is consistent with evidence that gustatory evaluation is strongly shaped by personal experience and cultural background (Allen et al., 2008), underscoring that individual differences provide a more accurate account than group-level comparisons alone.

Hunger had no main effect on pleasantness but selectively reduced pleasantness for low-concentration stimuli, consistent with the notion that hunger enhances wanting rather than liking (Rogers et al., 2021). Prior work shows that hunger effects on pleasantness are generally modest and depend on sensory richness (Khorisantono et al., 2024). The present data suggest that strong metabolic drive paired with weak gustatory input may create a mismatch between expectation and sensory payoff, reducing pleasantness. BMI showed no effects, likely due to the restricted, non-obese sample.

#### 4.1.2 Neural Taste Effects

The paradigm produced reliable activation in core gustatory pathways. Contrasting the taste stimuli with the rinses elicited strong activation in VPM, insula, postcentral gyrus, basal ganglia, and amygdala, reflecting sensory, motor, and affective components of taste delivery. Considering the contrast against Rinse, the unusually widespread activation was surprising, as motion-related activity like swallowing would not be anticipated. Amygdala engagement indicated that even simple taste stimuli undergo rapid salience and affective tagging (J. O’Doherty et al., 2001; Small et al., 2003). Left-sided activation is consistent with reports linking the left amygdala to pleasantness-related appraisal (Lanteaume et al., 2006), although hemispheric interpretations remain unstable across studies (Zald, 2003). Familiar and affectively clear stimuli tend to recruit the amygdala more reliably than ambiguous ones, which aligns with the stronger behavioral and neural responses to sweet relative to umami.

Insular activation was largely left-lateralized. Although hemispheric asymmetries in gustatory processing remain inconclusive (Dalenberg et al., 2015; Iannilli & Gudziol, 2019; Kurth et al., 2010), the current findings suggest a left-hemispheric bias for taste representation. The Stimulus > Rinse and High Concentration > Low Concentration contrasts elicited similar activation patterns, with corrected clusters mainly in the left anterior insula. While some studies have linked the left anterior insula to pleasantness coding (Dalenberg et al., 2015; Frank et al., 2008; Small et al., 2003), the current data show left-sided activation for taste presence and intensity, aligning with Iannilli & Gudziol (2018), who also acknowledged substantial interindividual variability and the absence of a fixed hemispheric dominance. The absence of right anterior insula activation contrasts with previous load-based studies (Duif et al., 2020; van Meer et al., 2023), potentially reflecting reduced stimulus complexity (compared to e.g. chocolate milk) and the inclusion of a neutral baseline, which may bias processing toward detection-oriented mechanisms. Overall, these results support a left-lateralized insular contribution to both taste detection and intensity coding, without implying fixed hemispheric dominance.

Taste category produced no differential activation in gustatory cortex, matching the lack of a behavioral effect of taste on intensity ratings. Both directions of the Sweet–Umami comparison involved fusiform and lateral occipital cortices and the angular gyrus, regions linked to semantic and cross-modal processes (Huerta et al., 2014; Kuhnke et al., 2023; Seghier, 2012). Sweet stimuli likely required little interpretive effort due to their strong familiarity and positive valence (Ventura & Mannella, 2011), whereas the unfamiliar MSG solution may have engaged additional conceptual evaluation to resolve its hedonic ambiguity.

Behavioral and neural data confirm that the paradigm reliably engaged gustatory pathways, with consistent left-lateralized insular and thalamic responses indicating sufficient sensitivity to detect taste effects. The absence of insular modulation in distraction contrasts thus likely reflects a genuine null result rather than insufficient statistical power.

### 4.2 Effects of Distraction on Taste Perception

#### 4.2.1 Behavioral Distraction Effects

Distraction reduced perceived intensity and pleasantness, consistent with prior evidence that attentional load dampens both sensory and hedonic components of taste processing (Ruda, Chellapandian, & Freiherr, 2024; van Meer et al., 2023). These effects occurred independently of taste category and concentration, indicating a global attenuation of taste perception rather than condition-specific modulation. While some prior studies reported stronger effects for high-concentration stimuli (Razzaghi-Asl et al., 2024; van der Wal & van Dillen, 2013; van Meer et al., 2023), the present naturalistic, video-based paradigm likely introduced more complex behavioral dynamics, leading to broader effects and reducing sensitivity to interactions.

#### 4.2.2 Neural Distraction Effects

High distraction elicited robust activation in bilateral occipital cortex, lingual gyrus, precuneus, and cerebellar lobules VIIIb/IX, reflecting enhanced engagement of visual and attention-related networks. These effects were consistent across taste and concentration levels, supporting the notion that visual distraction reallocates attentional resources toward external perceptual input, competing with internal gustatory processing (Lavie et al., 2004; M. G. Veldhuizen et al., 2012). VPM engagement further indicates early-stage competition between sensory modalities, consistent with salience-gating models of attentional prioritization (Wolff et al., 2021; Zhou et al., 2021). In contrast to typical cognitive load paradigms (van Dillen & van Steenbergen, 2018; van Meer et al., 2023), frontal activation was absent, suggesting that passive video viewing primarily induced bottom-up attentional capture rather than top-down executive control (Su et al., 2021).

Despite strong behavioral effects, canonical gustatory regions did not show significant attenuation. Preserved insular activity under high distraction suggests that attentional competition reduced perceptual salience without suppressing primary gustatory processing. Additionally, the insula’s role in multisensory attention likely limits its diagnostic utility in distraction paradigms; high intrinsic baseline activity and functional heterogeneity across sensory, attentional, and interoceptive domains (Bermudez-Rattoni, 2014; Chang et al., 2013; Gogolla, 2017) may mask small condition-specific effects, obscuring subtle modulations in group-level BOLD analyses. Null effects in OFC across contrasts might indicate that hedonic evaluation was not sufficiently recruited. Stimulus simplicity likely further reduced the need for OFC integration, consistent with prior evidence that OFC activation is most robust during explicit affective appraisal tasks (Small et al., 2007; van der Wal & van Dillen, 2013; Maria G. Veldhuizen & Small, 2011). The only regions showing reduced activation under high distraction were small bilateral clusters in the superior lateral occipital cortex or angular gyrus, which may reflect diminished integration of taste-related information into memory.

Notably, sweet stimuli under high distraction elicited bilateral amygdala activation compared to umami, suggesting that emotionally salient tastes can enhance affective processing and attentional engagement in parallel sensory networks (Zald, 2003). In contrast, high-concentration stimuli recruited additional thalamic and occipito-temporal regions, indicating that salient gustatory input increases demands on salience-gating mechanisms and attentional capacity.

ROI analyses corroborated the main pattern. Neither left anterior nor left middle insula showed significant distraction-related changes. Collectively, these results indicate that distraction induces a global reallocation of attentional resources, likely preserving primary gustatory representation but reducing its relative perceptual weight, a pattern consistent with resource-competition models rather than active inhibition.

### 4.3 Synthesis and Implications

The combined behavioral and neural findings reveal a dissociation between subjective taste experience and measurable neural modulation. While distraction significantly reduced perceived intensity and pleasantness, no corresponding attenuation of gustatory cortex activity was observed. This does not imply absent taste processing but suggests that gustatory signals remained subthreshold in group-level analyses, likely due to attentional competition from visually dominant input. Distraction thus reduced the perceptual impact of taste stimuli, which contributed less to conscious experience when attention was drawn elsewhere, reconciling the behavioral attenuation with the absent neural correlates.

Notably, the behavioral effects exceeded those of previous comparable fMRI studies (Duif et al., 2020; van Meer et al., 2023). This dissociation highlights a central challenge in neuroimaging research: Subjective changes can occur without corresponding BOLD differences due to interindividual variability, dynamic network compensation, or limited sensitivity of group statistics. The presence or absence of activation thus cannot be equated with the presence or absence of psychological effects.

Although speculative, the present dissociation between the extent of intensity (d = –0.28) and pleasantness (d = –0.21) suppression may offer a mechanistic explanation: Reduced intensity likely reflects diminished perceptual vividness and may weaken satiety signaling, while preserved pleasantness sustains the subjective reward value of food and supports continued consumption. In other words, food may feel “less vivid” but still “good enough” to maintain intake under diminished sensory awareness. However, this difference was modest and should not be taken as definitive evidence for dissociable mechanisms. Pleasantness was influenced by multiple interacting factors in the behavioral model and showed greater interindividual variability, which may explain both its relative robustness to distraction and the absence of consistent OFC activation at the group level.

The absence of OFC attenuation further suggests that the hedonic value of food was not substantially diminished by distraction. As the OFC plays a key role in representing the affective significance of taste (Bender et al., 2009; Duif et al., 2020; Small et al., 2007), a decrease in OFC activation would likely have been detected if distraction had meaningfully reduced hedonic evaluation.

Consequently, distraction effects should not be interpreted solely based on suppression of gustatory regions. Our results align with prior work showing intact insular activity but disrupted connectivity with attentional and reward-related networks (Duif et al., 2020). Bidirectional dynamics may contribute. Strong gustatory stimulation may occupy insular–thalamic bandwidth, reducing attentional processing of external visual input (Menon, 2025), whereas weak or ambiguous tastes may recruit compensatory attentional processing, potentially explaining differential occipital and parietal engagement between sweet and umami stimuli.

Only a few prior studies have explicitly examined hedonic aspects of taste under distraction, but those that did reported modest reductions in perceived pleasantness (Ruda, Chellapandian, & Freiherr, 2024; van Rijn et al., 2018). This convergence suggests that hedonic evaluation, while more resilient than sensory discrimination, remains susceptible to attentional modulation. This may partially explain why distracted eating can increase food intake, even when taste intensity is reduced.

Taken together, these findings indicate that distraction alters how, not whether taste is processed. Neural resources are redistributed within visual–attentional networks, reducing perceptual salience and reweighting taste signals while preserving primary gustatory processing, explaining why eating under distraction diminishes intensity yet maintains hedonic properties.

### 4.4 Methodological Considerations, Strengths, and Limitations

The videos presented were generic and non-individualized, whereas real-life distractions are often personalized, aligning with individual preferences and emotional states (Su et al., 2021). Contemporary short-form media exerts even stronger attentional capture (Chiencharoenthanakij et al., 2025; Uncapher & Wagner, 2018; Yan et al., 2024), suggesting that our neural results might represent a conservative estimate of real-world effects. Full-length films could enhance naturalism at the cost of temporal control; future studies might consider such paradigms without in-scanner ratings to maximize immersion. Similarly, the use of simple single-compound taste stimuli limited the ability to probe neural sensitivity to more complex or ecologically valid flavors. While MSG acted as a known flavor enhancer, it may have introduced additional perceptual or neural variance. The binary taste design ensured experimental control but constrained ecological validity, potentially contributing to bimodal intensity ratings and limiting mid-range interpretability. Residual taste effects from the rinse baseline may also have introduced minor variance, emphasizing the value of explicit baseline ratings in future studies. These considerations highlight the need for future studies to integrate parametric measures of attention — such as eye tracking, pupillometry, or in-task ratings — with personalized visual and complex gustatory stimuli.

Most participants reported frequent engagement with digital media during meals, and the passive visual stimuli effectively competed for attention without invoking explicit executive control, as evidenced by strong visual and thalamic activation but absent prefrontal recruitment. While the implemented conditions addressed the main aim of comparing attention toward versus away from food, the absence of a truly neutral baseline limits the interpretability of the low-distraction condition. Including a no-visual-stimulation condition could allow a more continuous assessment of attentional load.

Distraction validation relied on retrospective ratings of subjective interest. Although differences were large (d > 1), these measures are vulnerable to memory bias and lack temporal resolution. Repeated in-task ratings could improve temporal resolution and enable parametric modulation, which was not feasible here because each block provided only one interest value, resulting in no within-regressor variability. Behavioral modeling of intensity ratings showed only moderate fit, particularly in the midrange, reflecting the binary stimulus design and interpretational ambiguities of the VAS, especially if the midpoint was implicitly interpreted as neutral.

Neural measurements may have been influenced by technical constraints. The gustometer allowed precise event-related taste delivery, but swallowing-induced motion introduced residual artifacts, particularly in frontal regions. Continuous-flow systems (Kami et al., 2008) or external motion monitoring (e.g., EMG) could mitigate these effects. While the current paradigm cannot fully emulate complex meals, adaptations for more viscous or particulate stimuli could enhance ecological validity and allow exploration of additional taste dimensions such as fat content.

## 5 Conclusion and Future Directions

Distraction modulated taste perception not by suppressing gustatory cortex activity but by redistributing processing resources, reducing the relative salience of taste. Behaviorally, perceived intensity was attenuated more than pleasantness, indicating sensory vividness is more vulnerable to attentional interference than hedonic evaluation. Neural responses in canonical gustatory regions (insula, OFC) were not significantly modulated; instead, additional activation was observed in occipital, thalamic, and cerebellar regions, reflecting bottom-up capture by visual input. These findings support a resource-competition model in which taste signals remain represented but are assigned reduced cognitive weight.

The study demonstrates that realistic, video-based paradigms can be implemented in fMRI without sacrificing experimental precision, bridging laboratory tasks and real-world eating contexts. Distraction reliably alters subjective taste experience, emphasizing that susceptibility to distracted eating varies with individual attentional engagement and gustatory sensitivity.

Future work should employ parametric, personalized distraction stimuli (e.g., self-selected videos or social-media content) combined with concurrent engagement measures. Complex, multisensory taste stimuli could clarify whether attentional interference affects early sensory processing, integration, or memory formation.

Reduced taste salience under distraction may weaken satiety signaling while preserving hedonic value, potentially promoting overeating. Understanding this mechanism could inform interventions fostering mindful eating, maintaining both healthy consumption and the motivational quality of eating in media-rich environments.

### Data and code availability statement

Due to ethical restrictions concerning participant confidentiality, the dataset supporting this study is available from the corresponding author upon reasonable request and subject to institutional approval.

## Acknowledgements

This work was supported by the Brain Imaging Facility of the Interdisciplinary Center for Clinical Research (IZKF) Aachen, within the Faculty of Medicine at RWTH Aachen University. This project was funded by the K1 Commission of FH Aachen University of Applied Sciences. We would like to thank all the participants for their contribution to the study, as well as the volunteers who took part in the pilot measurements. We also thank CoasterForce and Star Wars Collateral Story on YouTube for granting permission to use their videos.

## CRediT authorship contribution statement

**Robert Friedmann:** Conceptualization, Methodology, Software, Investigation, Formal analysis, Funding acquisition, Writing – original draft. **Bichr Grii:** Methodology, Investigation. **Esther Jacoby:** Methodology, Investigation. **Ilya Digel:** Funding acquisition, Writing – review and editing. **Rea Rodriguez-Raecke:** Conceptualization, Writing – review and editing, Resources. **Rik Sijben:** Conceptualization, Investigation, Supervision, Writing – review and editing

## Declaration of Generative AI

During the preparation of this work, ChatGPT was used to assist with phrasing and text optimization. The authors reviewed and edited the content as needed and take full responsibility for the content of the published article.

