## Supplementary Materials for "Engaging Visual Media Shifts Taste-Related Neural Processing: An fMRI Study on Distracted Eating"

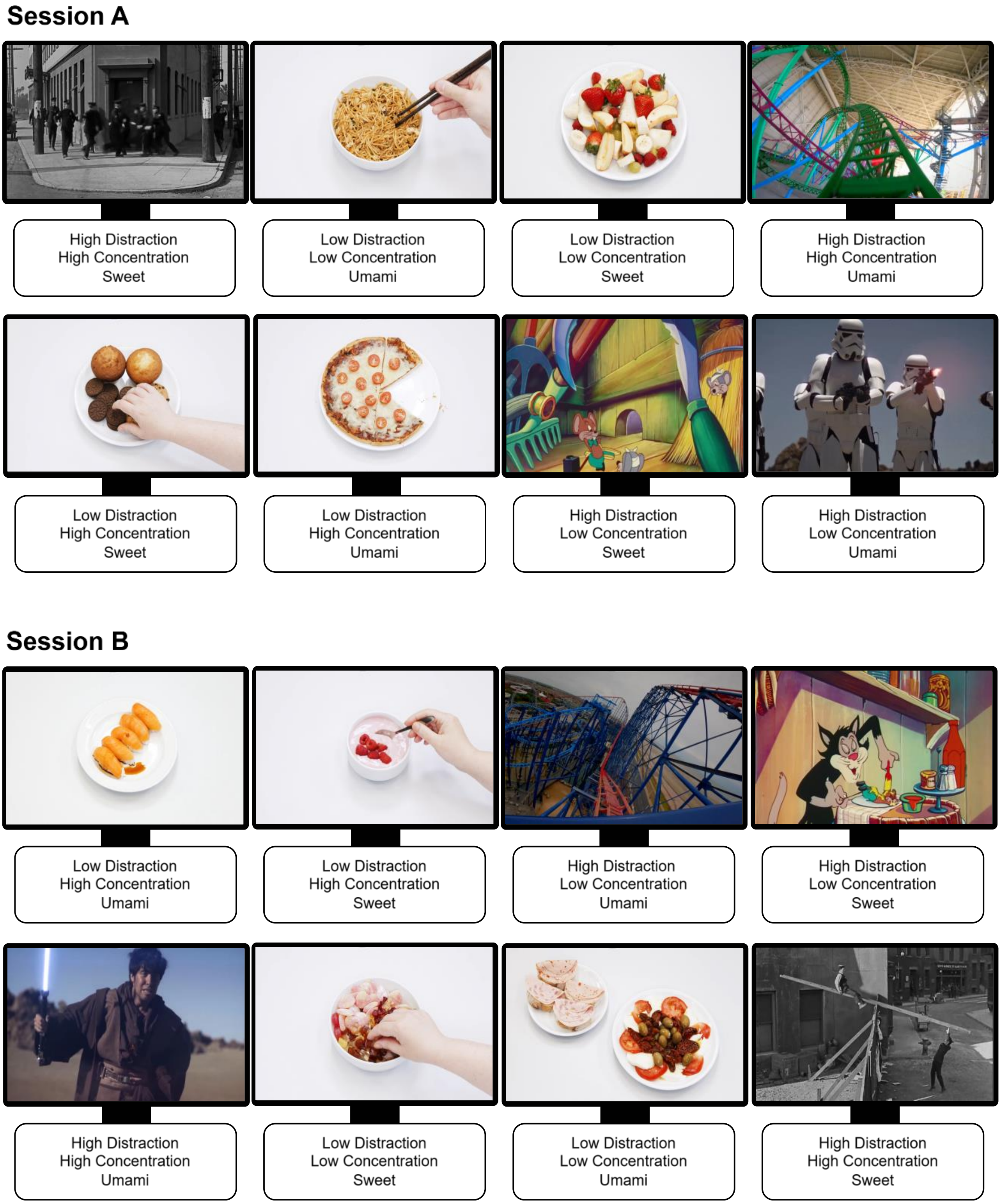

*Supplementary Figure 1: Depiction of the exact block order as used in the experiment. Block order was consistent across participants. Both experimental sessions covered all conditions of the 2×2×2 factorial within-subject design. Food-related videos used as low distraction were filmed at the Audiovisual Media Center of the Medical Faculty of RWTH Aachen University. Film-related videos used as high distraction were public domain or used with the creators' permission (Chaplin, 1928; CoasterForce, 2023, 2024; Jones, 1939; Keaton, 1922; Star Wars Collateral Story, 2023). Full source information is provided in the references.*

Levene's tests and a Fligner-Killeen tests confirmed homoscedasticity across sexes for Intensity Rating (Levene's $F(1, 631) = 0.0007$, $p = 0.979$; Fligner-Killeen $\chi^2(1) = 0.08$, $p = 0.777$), whereas for Pleasantness Rating, both tests revealed significant heterogeneity of variance (Levene's $F(1, 631) = 34.31$, $p < 0.001$; Fligner-Killeen $\chi^2(1) = 30.01$, $p < 0.001$). Based on these indications, Sex was included as a fixed effect in the subsequent models predicting Pleasantness Rating.

Intensity Rating and Pleasantness Rating were plotted across levels of Concentration and Taste, connecting individual participants to visualize within-subject changes (Supplementary Figure 2). The plots revealed strong, uneven individual effects of Taste, but not Concentration, leading to the inclusion of a random slope for Taste in the subsequent models predicting both Intensity Rating and Pleasantness Rating.

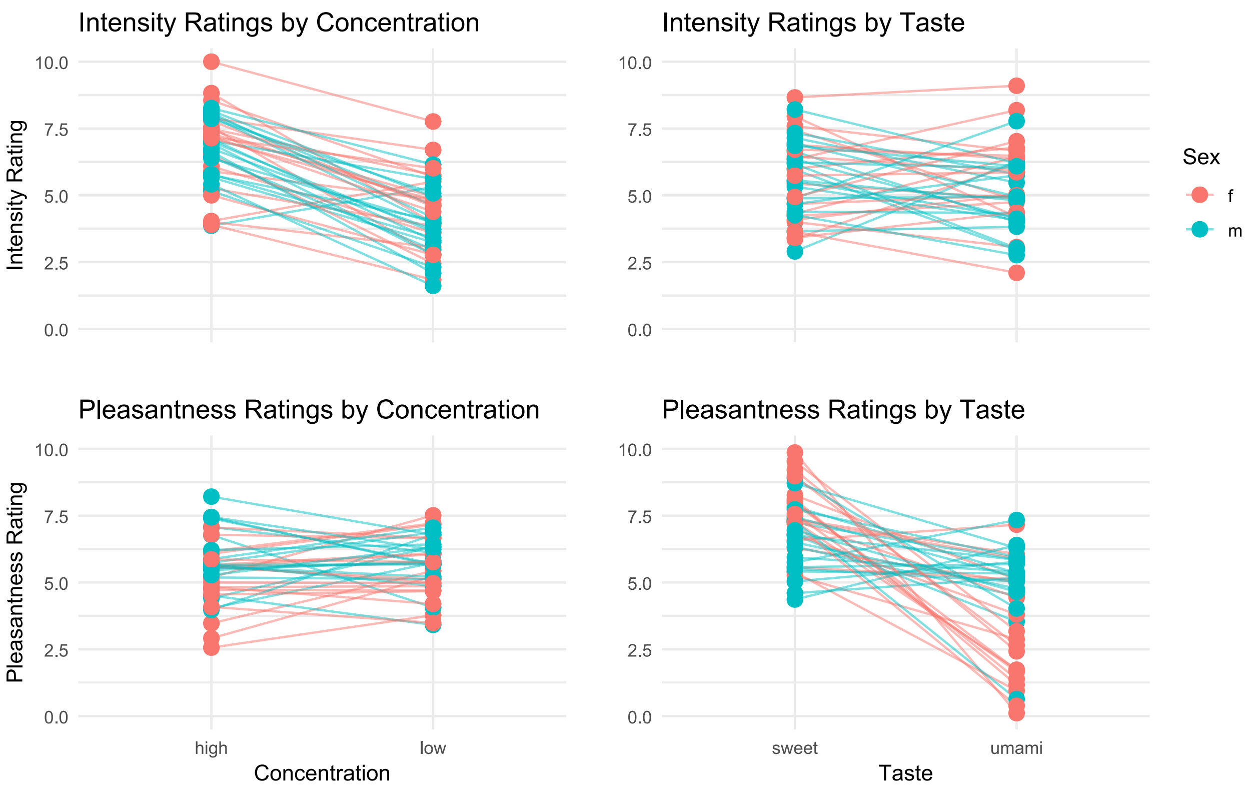

*Supplementary Figure 2: Intensity Rating and Pleasantness Rating plotted for all levels of Concentration and Taste, connecting individual participants to visualize inter-group and interindividual variability, with male participants depicted in blue and female participants in pink.*

The differences between levels of Concentration were tested both within and between tastes to assess distinctiveness. Paired t-tests comparing Intensity Rating yielded a significant difference between high and low Concentration within both tastes (both $p < 0.001$) but not across tastes ($p_{high} = 0.465$, $p_{low} = 0.237$), indicating that taste categories were matched in baseline intensity. This confirmed that the stimulus design successfully manipulated perceived intensity while maintaining comparability between the two tastes (Supplementary Figure 3).

The effectiveness of the distraction manipulation was verified using the subjective Interest Rating. A paired t-test comparing Interest Rating between the levels of Distraction revealed a significant difference with a large effect size ($p < 0.001$, $d = 1.31$). Film-related videos were on average rated as more interesting than food-related videos, confirming that the two conditions successfully induced different levels of attentional engagement (Supplementary Figure 3).

Paired t-tests for Intensity Rating, Pleasantness Rating and Interest Rating across sessions revealed no significant differences (all $p > 0.1$). Therefore, session was not included as a factor in the behavioral models.

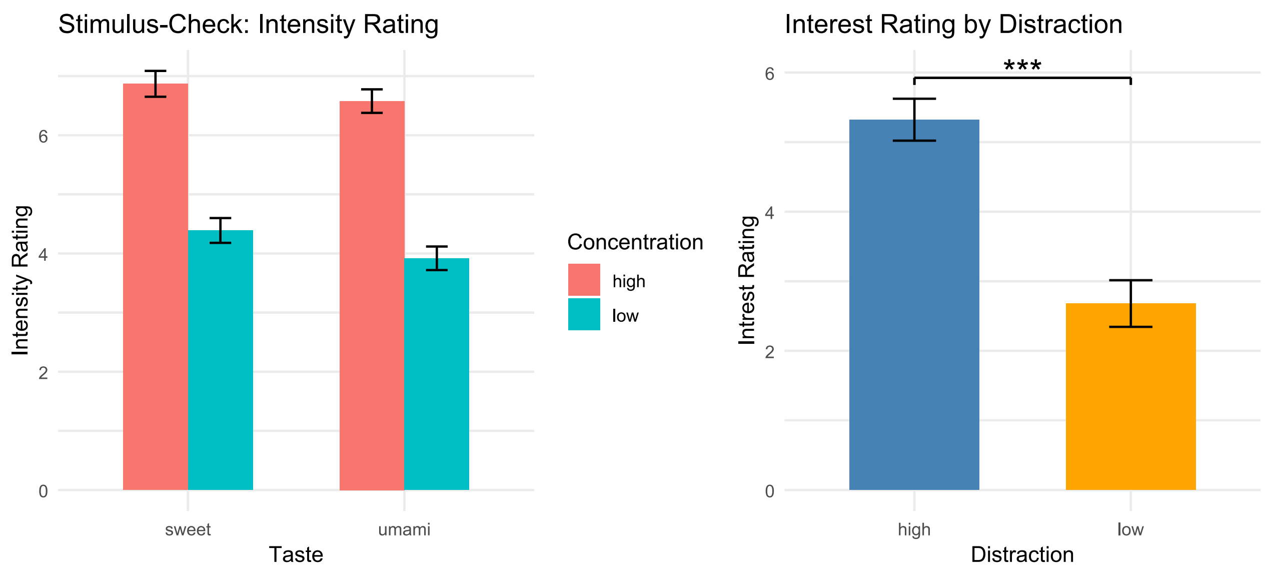

*Supplementary Figure 3: (Left) Average Intensity Rating for both concentrations and tastes, confirming distinctiveness between concentrations and comparability between tastes. (Right) Average Interest Rating for high and low Distraction, indicating a successfully induced variability in attentional engagement. Vertical lines represent standard error. *** indicates p < 0.001*

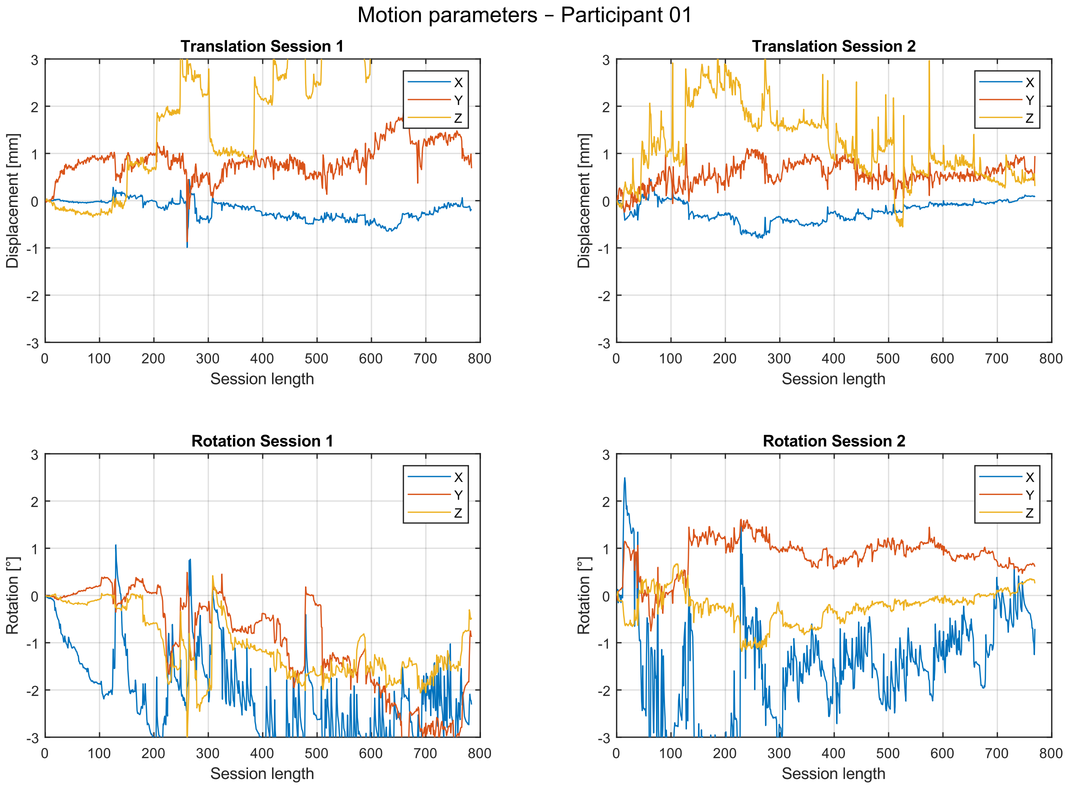

*Supplementary Figure 4: Exemplary motion parameters for all 6 spatial degrees of freedom for both scanning sessions. Prominent displacement of > 2 mm or rotation of > 2° was deemed problematic and led to the exclusion of two participants from the fMRI analysis.*

| Session A | | | | | | | | | | |
|---|---|---|---|---|---|---|---|---|---|---|
| **Video 1** | H S | H S | H S | H S | H U | H S | L U | H S | H S | H S |
| **Video 2** | L U | L U | L U | L U | L U | H S | L U | L U | L S | L U |
| **Video 3** | L S | L S | H U | L S | L S | L S | L S | L U | L S | L S |
| **Video 4** | H U | H U | H U | L S | H U | H S | H U | H U | H U | H U |
| **Video 5** | H S | H S | H S | H S | H U | H S | L U | H S | H S | H S |
| **Video 6** | H U | H U | H U | L S | H U | H S | H U | H U | H U | H U |
| **Video 7** | L S | L S | H U | L S | L S | L S | L S | L U | L S | L S |
| **Video 8** | L U | L U | L U | L U | L U | H S | L U | L U | L S | L U |

| Session B | | | | | | | | | | |
|---|---|---|---|---|---|---|---|---|---|---|
| **Video 1** | H U | H U | H U | H U | H S | H U | H U | L S | H U | H U |
| **Video 2** | H S | H S | H S | H S | H U | H S | L U | H S | H S | H S |
| **Video 3** | L U | L U | L S | L U | L U | H S | L U | L U | L U | L U |
| **Video 4** | L S | L S | L U | L S | L S | L S | H U | L S | L S | L S |
| **Video 5** | H U | H U | H U | H U | H S | H U | H U | L S | H U | H U |
| **Video 6** | L S | L S | H U | L S | L S | L S | L U | L S | L S | L S |
| **Video 7** | L U | L U | L S | L U | L U | H S | L U | L U | L U | L U |
| **Video 8** | H S | H S | H S | H S | H U | H S | L U | H S | H S | H S |

*Supplementary Table 1: Depiction of the exact stimulus order as used in the experiment. One taste stimulus was delivered every 15 seconds, followed by a rinse 5 seconds later. H = High Concentration, L = Low Concentration, S = Sweet, U = Umami. Each block comprised 8 target stimuli and 2 distractor stimuli. Distractor stimuli used to avoid habituation are highlighted in red.*

| Label | Side | MNI coordinates [mm] | No. of voxels | p-value | T-value (peak) |
|---|---|---|---|---|---|
| **(A) Stimulus > Rinse** | | | | | |
| Inferior lateral occipital, middle temporal, posterior cingulate, anterior cingulate, precuneus, superior parietal lobule, postcentral, thalamus, putamen, caudate, posterior insula, lingual, cerebellum IX, cerebellum Crus II, brainstem | L+R | -44 -64 0 | 58403 | < 0.001 | 10.68 |
| ↳ Precuneus | L+R | 8 -50 50 | – | < 0.001 | 9.49 |
| Inferior frontal, pars opercularis | R | 58 28 22 | 997 | < 0.001 | 7.71 |
| Anterior temporal fusiform, amygdala | R | 34 -6 -32 | 222 | < 0.001 | 7.53 |
| Amygdala, anterior temporal fusiform | L | -30 2 -24 | 223 | 0.001 | 6.76 |
| ↳ Anterior temporal fusiform | L | -38 -6 -34 | – | 0.004 | 6.18 |
| Inferior frontal, pars opercularis | L | -42 8 24 | 414 | 0.002 | 6.43 |
| ↳ Amygdala | R | 22 -4 -16 | – | 0.012 | 5.80 |
| Anterior insula | L | -30 20 6 | 88 | 0.007 | 6.01 |
| Precentral | L | -28 -8 60 | 91 | 0.014 | 5.74 |
| Hippocampus | R | 22 -16 -22 | 4 | 0.021 | 5.57 |
| Middle insula | L | -38 -4 12 | 10 | 0.023 | 5.53 |
| Frontal medial | | 0 52 -10 | 21 | 0.022 | 5.55 |
| Middle insula | R | 36 2 12 | 4 | 0.035 | 5.36 |

**(A) Rinse > Stimulus**

| | | | | | |
|---|---|---|---|---|---|
| Ventricles (artefact) | – | – | – | – | – |

**(B) High Concentration > Low Concentration** *($p_{FWEc} < 0.05$)

| | | | | |
|---|---|---|---|---|
| Occipital pole, intracalcarine, lingual, precuneus | R | 10 -86 6 | 11152 | < 0.001 |
| ↳ Occipital pole | L | -6 -90 2 | – | < 0.001 |
| Anterior Insula, middle insula, thalamus, postcentral | L | -30 8 10 | 5909 | < 0.001 |
| Anterior cingulate | L | -6 6 36 | 233 | 0.032 |
| Postcentral gyrus, anterior insula, middle insula, thalamus | R | 64 2 22 | 4068 | < 0.001 |
| Juxtapositional lobule | | 2 -6 76 | 1029 | < 0.001 |
| Posterior cingulate | R | 4 -38 20 | 231 | 0.033 |

**(B) Low Concentration > High Concentration** *($p_{FWEc} < 0.05$)

| | | | | |
|---|---|---|---|---|
| – | – | – | – | – |

**(C) Sweet > Umami**

| | | | | | |
|---|---|---|---|---|---|
| Inferior lateral occipital | L | -52 -74 6 | 77 | < 0.001 | 8.33 |
| Angular | R | 52 -52 18 | 279 | 0.003 | 6.57 |
| Inferior lateral occipital | R | 56 -62 6 | 42 | 0.016 | 5.97 |
| Temporal occipital fusiform | R | 44 -48 -20 | 5 | 0.028 | 5.74 |
| Temporal occipital fusiform | L | -42 -50 -22 | 2 | 0.042 | 5.56 |

**(C) Umami > Sweet**

| | | | | | |
|---|---|---|---|---|---|
| Temporal occipital fusiform | R | 28 -48 -10 | 416 | < 0.001 | 10.12 |
| Superior lateral occipital | R | 32 -76 18 | 1345 | < 0.001 | 9.67 |
| Superior lateral occipital | L | -20 -80 26 | 705 | < 0.001 | 7.56 |
| Superior lateral occipital | L | -22 -64 62 | 133 | 0.001 | 7.34 |
| Lingual | L | -26 -46 -8 | 225 | 0.001 | 6.98 |
| Superior lateral occipital | R | 20 -74 46 | 57 | 0.006 | 6.32 |
| Superior lateral occipital | R | 22 -60 52 | 45 | 0.007 | 6.27 |

**(D) High Distraction > Low Distraction**

| | | | | | |
|---|---|---|---|---|---|
| Superior lateral occipital, inferior lateral occipital, supracalcarine, intracalcarine, lingual, temporal occipital fusiform, R thalamus | L+R | -10 -94 12 | 19440 | < 0.001 | 19.57 |
| Cerebellum VIIIb / IX | L | -14 -48 -50 | 309 | < 0.001 | 10.48 |
| Thalamus | L | -18 -30 0 | 178 | < 0.001 | 8.73 |
| Cerebellum VIIIb / IX | R | 16 -50 -50 | 163 | < 0.001 | 8.12 |
| Cerebellum Vermis IX | | 0 -60 -40 | 22 | 0.004 | 6.74 |
| Superior frontal gyrus | R | 28 0 58 | 27 | 0.004 | 6.70 |
| Cerebellum VIIb | L | -8 -76 -44 | 22 | 0.004 | 6.66 |
| Superior temporal gyrus, posterior division | R | 52 -30 2 | 11 | 0.006 | 6.65 |

| | | | | | |
|---|---|---|---|---|---|
| **(D) Low Distraction > High Distraction** | | | | | |
| Ventricles (artefact) | – | – | – | – | – |
| Superior lateral occipital / Angular | L | -50 -62 48 | 22 | 0.005 | 6.59 |
| Superior lateral occipital / Angular | R | 48 -56 54 | 31 | 0.007 | 6.47 |

*Supplementary Table 2: Summary of brain regions exhibiting main effects of (A) Stimulus, (B) Concentration, (C) Taste and (D) Distraction. A, C, D significant at $p_{FWE}$ < 0.05; B significant at $p_{FWEc}$ < 0.05 to improve sensitivity. Region labels with a slash (/) denote spatially overlapping or not clearly separable clusters based on the applied anatomical atlas. ↳ denotes sub-peaks within the same cluster. Anatomical labels were assigned using the Harvard–Oxford cortical and subcortical atlases; cerebellar regions were labeled using the probabilistic cerebellar atlas (MNI-152); the amygdala was defined using the Jülich histological atlas.*

| Label | Side | MNI coordinates [mm] | No. of voxels | p-value | T-value (peak) |
|---|---|---|---|---|---|
| **(A) Interaction: Distraction (High > Low) × Concentration (High > Low)** *($p_{FWEc}$ < 0.05) | | | | | |
| Occipital pole, intracalcarine | L | -12 -92 4 | 3096 | < 0.001 | |
| **(A) Interaction: Distraction (Low > High) × Concentration (High > Low)** *($p_{FWEc}$ < 0.05) | | | | | |
| Inferior temporal, inferior lateral occipital | R | 48 -52 -10 | 2869 | < 0.001 | |
| Inferior lateral occipital | L | -46 -78 -8 | 2329 | 0.001 | |
| Superior parietal lobule | R | 34 -40 72 | 227 | 0.032 | |
| **(B) Interaction: Distraction (High > Low) × Taste (Sweet > Umami)** *($p_{FWEc}$ < 0.05) | | | | | |
| Inferior lateral occipital, middle temporal, inferior temporal, temporal occipital fusiform, postcentral, superior parietal lobule | R | 46 -64 4 | 9345 | < 0.001 | |
| Inferior lateral occipital, middle temporal, inferior temporal, temporal occipital fusiform, angular | L | -48 -68 2 | 4708 | < 0.001 | |
| Postcentral, superior parietal lobule | L | -46 -34 52 | 1844 | 0.001 | |
| Anterior temporal fusiform | R | 36 -8 -34 | 459 | 0.002 | |
| Middle frontal / Inferior frontal, pars opercularis | R | 52 18 32 | 1270 | < 0.001 | |
| Thalamus | L | -16 -28 8 | 317 | 0.011 | |
| **(B) Interaction: Distraction (Low > High) × Taste (Sweet > Umami)** *($p_{FWEc}$ < 0.05) | | | | | |
| Ventricles (artefact) | – | – | – | – | |

*Supplementary Table 3: Summary of brain regions exhibiting interaction effects of (A) Distraction × Concentration and (B) Distraction × Taste. All significant at $p_{FWEc}$ < 0.05 to improve sensitivity. Region labels with a slash (/) denote spatially overlapping or not clearly separable clusters based on the applied anatomical atlas. Anatomical labels were assigned using the Harvard–Oxford cortical and subcortical atlases; cerebellar regions were labeled using the probabilistic cerebellar atlas (MNI-152).*

| Label | Side | MNI coordinates [mm] | No. of voxels | p-value | T-value (peak) |
|---|---|---|---|---|---|
| **(A) [High Distraction > Low Distraction]$_{HighConcentration}$** | | | | | |
| Occipital pole, intracalcarine, lingual | L | -12 -94 12 | 9879 | < 0.001 | 21.77 |
| Cerebellum VIIIb / IX | L | -14 -50 -50 | 105 | < 0.001 | 8.17 |
| Precuneus | R | 6 -54 58 | 354 | 0.001 | 7.13 |
| Precuneus | L | -6 -46 60 | 131 | 0.002 | 7.06 |
| Cerebellum VIIIb / IX | R | 16 -48 -52 | 47 | 0.002 | 6.90 |
| Thalamus | R | 20 -28 -2 | 4 | 0.005 | 6.58 |
| Thalamus | L | -24 -30 0 | 28 | 0.006 | 6.54 |
| Temporal occipital fusiform | R | 42 -42 -24 | 24 | 0.010 | 6.34 |
| Cerebellum Crus II | | -2 -74 -32 | 2 | 0.024 | 5.98 |
| Angular | R | 52 -44 14 | 26 | 0.029 | 5.76 |
| **(A) [Low Distraction > High Distraction]$_{HighConcentration}$** | | | | | |
| – | – | – | – | – | – |
| **(B) [High Distraction > Low Distraction]$_{LowConcentration}$** | | | | | |
| Inferior lateral occipital, temporal occipital fusiform, lingual | L+R | 48 -74 6 | 15009 | < 0.001 | 16.03 |
| Cerebellum VIIIb / IX | L | -14 -48 -50 | 150 | < 0.001 | 9.05 |
| Precuneus | R | 8 -46 62 | 478 | < 0.001 | 8.14 |
| Posterior supramarginal / Middle temporal | R | 50 -40 12 | 228 | < 0.001 | 8.01 |
| Cerebellum VIIIb / IX | R | 16 -50 -50 | 62 | 0.001 | 7.13 |
| Cerebellum VIIb | L | -8 -76 -44 | 3 | 0.018 | 6.10 |
| Precuneus | L | -10 -46 66 | 6 | 0.040 | 5.78 |
| **(B) [Low Distraction > High Distraction]$_{LowConcentration}$** | | | | | |
| Ventricles (artefact) | – | – | – | – | – |
| **(C) [High Distraction > Low Distraction]$_{Sweet}$** | | | | | |
| Inferior lateral occipital, intracalcarine, lingual | R | 46 -66 4 | 17640 | < 0.001 | 15.70 |
| Thalamus | L | - 18 – 28 6 | 256 | < 0.001 | 8.73 |
| Thalamus | R | 18 -28 0 | 175 | < 0.001 | 8.35 |
| Amygdala laterobasal group, anterior temporal fusiform | R | 24 -6 -24 | 226 | < 0.001 | 8.35 |
| Cerebellum VIIIb | L | -12 -50 -52 | 157 | < 0.001 | 7.77 |
| Cerebellum VIIIb | R | 26 -38 -46 | 91 | < 0.001 | 7.63 |
| Temporal pole | R | 50 14 -30 | 80 | 0.001 | 7.54 |
| Cerebellum VIIb / Crus II | L | -8 -76 -44 | 33 | 0.002 | 6.91 |
| Superior lateral occipital | L | -22 -72 36 | 27 | 0.003 | 6.77 |
| Precentral | R | 44 6 30 | 71 | 0.004 | 6.66 |
| Amygdala laterobasal group, temporal pole | L | -28 0 -24 | 29 | 0.004 | 6.65 |
| ↳ Temporal pole | L | -26 6 -30 | – | 0.010 | 6.31 |
| Anterior middle temporal / Anterior superior temporal | R | 52 -2 -22 | 20 | 0.007 | 6.44 |
| Postcentral | R | 42 -32 56 | 18 | 0.013 | 6.19 |
| Postcentral | R | 36 -30 46 | 8 | 0.020 | 6.04 |
| Anterior supramarginal | R | 60 -32 32 | 4 | 0.030 | 5.86 |

**(C) [Low Distraction > High Distraction]$_{Sweet}$**

| | | | | | |
|---|---|---|---|---|---|
| Ventricles (artefact) | – | – | – | – | – |

**(D) [High Distraction > Low Distraction]$_{Umami}$**

| | | | | | |
|---|---|---|---|---|---|
| Lingual | R | 26 -42 -8 | 7977 | < 0.001 | 13.29 |
| Cerebellum VIIIb / IX | L | -14 -50 -50 | 86 | < 0.001 | 8.09 |
| Cerebellum VIIIb / IX | R | 16 -50 -50 | 17 | 0.005 | 6.62 |
| Postcentral /Precuneus | R | 8 -42 60 | 80 | 0.006 | 6.54 |
| Lateral occipital | L | -40 -76 18 | 99 | 0.007 | 6.46 |
| Postcentral /Precuneus | L | -14 -42 56 | 23 | 0.010 | 6.31 |
| Lingual | L | -20 -54 4 | 5 | 0.027 | 5.92 |

**(D) [Low Distraction > High Distraction]$_{Umami}$**

| | | | | | |
|---|---|---|---|---|---|
| Superior parietal lobule | L | -44 -42 56 | 116 | 0.006 | 6.56 |
| Angular / Posterior supramarginal | R | 50 -52 54 | 2 | 0.034 | 5.84 |

*Supplementary Table 4: Summary of brain regions exhibiting distraction effects at (A) High Concentration , (B) Low Concentration, (C) Sweet Taste and (D) Umami Taste. All significant at $p_{FWE} < 0.05$. Region labels with a slash (/) denote spatially overlapping or not clearly separable clusters based on the applied anatomical atlas. ↳ denotes sub-peaks within the same cluster. Anatomical labels were assigned using the Harvard–Oxford cortical and subcortical atlases; cerebellar regions were labeled using the probabilistic cerebellar atlas (MNI-152); the amygdala was defined using the Jülich histological atlas.*
